# Linking Oxygen-Induced Oxidative Stress to Resource Recovery by Enhancing the Production of Extracellular Polymeric Substances in Activated Sludge Microbial Communities

**DOI:** 10.1101/2025.04.29.651171

**Authors:** Xueyang Zhou, Bharat Manna, Boyu Lyu, Naresh Singhal

## Abstract

As the global transition toward circular wastewater treatment intensifies, extracellular polymeric substances (EPS) have emerged as valuable targets for resource recovery. Although most efforts have focused on aerobic granular sludge, conventional activated sludge systems, which account for most global wastewater treatment, remain underexploited for EPS recovery. Building on the established link between oxidative stress and EPS biosynthesis in pure strains, we propose that strategically manipulating oxygen exposure patterns to intensify oxidative stress in activated sludge microbial communities could enhance EPS production. To test this, we applied continuous oxygen perturbation under aerobic exposure to intensify oxidative stress. Compared to a stable oxygen condition simulating typical wastewater aeration, the perturbation considerably enhanced EPS yield to 74.4□mg/L/day, a 90.5% increase over the stable condition (39.0□mg/L/day). To validate the role of oxidative stress in EPS enhancement, intermittent anoxic phases were introduced into the perturbation pattern to relieve oxidative stress, causing the EPS-enhancing effect to disappear, with yield dropping to 9.8□mg/L/day. Mechanistically, intensified oxidative stress under aerobic continuous perturbation was primarily driven by elevated reducing substrates for non-respiratory flavoenzymes, exemplified by glutamate synthase, glutathione reductase, and dihydrolipoamide dehydrogenase, which are prone to generate H_2_O_2_ as an unintended metabolic byproduct. Among the multiple microbial groups contributing to H_2_O_2_ production, *Methylophilaceae, Comamonadaceae*, and *Rhodobacteraceae* were distinguished by simultaneously exhibiting upregulation of EPS biosynthesis proteins, suggesting that taxa within these families collectively mediated both H_2_O_2_ production and EPS enhancement. By modulating aeration, this study offers a chemical-free, controllable strategy for enhancing EPS production within conventional activated sludge systems.

**Graphical abstract:** 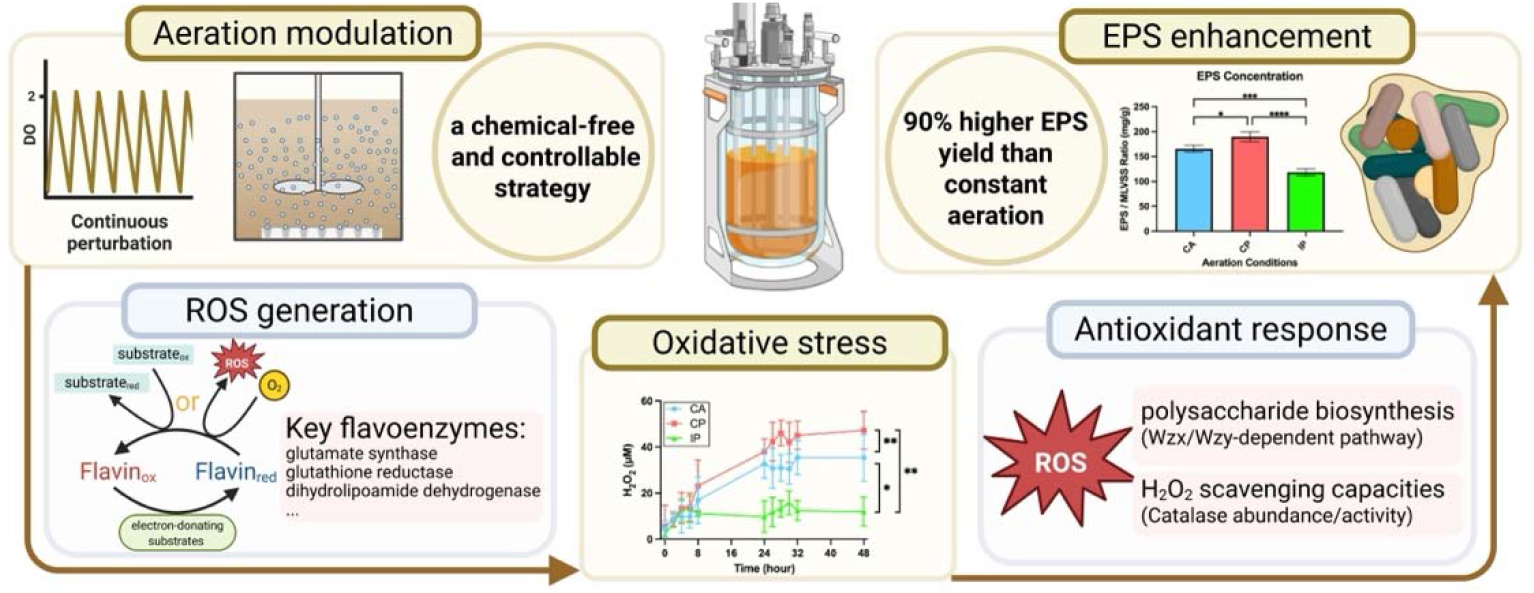

## 1. Introduction

Resource recovery from wastewater treatment plants (WWTPs) has gained increasing importance due to both economic incentives and environmental sustainability goals (Faragò et al., 2021). Among the various recoverable resources, extracellular polymeric substances (EPS) have attracted growing attention owing to their diverse functionalities and commercial value (Fang et al., 2024; Salama et al., 2016). In the Netherlands, the first two demonstration-scale projects have successfully implemented the commercial extraction of EPS from aerobic granular sludge, marketed under the trade name Kaumera (Bahgat et al., 2023). These extracted EPS are commercially viable and contribute substantially to the economic returns of resource recovery programs, with Kaumera potentially accounting for up to 50% of the turnover for the Energy and Raw Materials Factory (Bahgat et al., 2023). However, although EPS recovery has been successfully demonstrated in aerobic granular sludge systems, the majority of WWTPs worldwide still rely on conventional activated sludge (CAS) systems, which generate significantly larger volumes of sludge but offer limited feasibility for EPS recovery due to difficulties in enhancing EPS content within the biomass (Hao et al., 2019; Kroiss, 2004). Enhancing EPS yields and improving EPS quality in CAS systems therefore remains a key challenge and an important frontier in the development of sustainable wastewater biorefineries (Li et al., 2021, 2015; van Loosdrecht and Brdjanovic, 2014).

A range of strategies has been explored to enhance microbial EPS production, including external additives (e.g., powder activated carbon, zeolite, coagulants) (Ji et al., 2010; Yuniarto et al., 2013), exogenous quorum sensing signals (e.g., C14-AHLs, AHL cocktails) (González et al., 2013; Tan et al., 2014), nutrient optimization (Ji et al., 2010; Li and Yang, 2007; Ye et al., 2011), and environmental stress induction (such as salinity, pH shifts, and metal ions) (Shi et al., 2013; Shu and Lung, 2004). While these approaches can stimulate EPS biosynthesis, they often suffer from drawbacks such as high chemical costs, limited controllability, inconsistent performance, or environmental concerns (Shi et al., 2017). This highlights the need for a more cost-effective and operationally manageable strategy. Mechanistically, microorganisms naturally produce and secrete EPS in response to environmental stress as a protective strategy, facilitating cell aggregation and structural stability in biofilms or sludge flocs (Wingender et al., 1999). This inherent stress-response mechanism offers a potential oxidative stress-based approach to promote EPS production, which has predominantly been explored in pure cultures (Ding et al., 2024; Ionescu and Belkin, 2009). Intracellular reactive oxygen species (ROS) generated through endogenous metabolism can activate transcriptional regulators such as OxyR and influence other stress-response regulatory systems, including those involving OmpR, thereby promoting EPS biosynthesis (Seo et al., 2017; Jang et al., 2016).

Nevertheless, it remains unclear how complex mixed microbial communities respond under such conditions. Importantly, our recent observations revealed that specific oxygen perturbation regimes can induce intensified endogenous oxidative stress in activated sludge systems (Zhou et al., 2025), which may serve as a potential breakthrough toward enhancing EPS production in mixed microbial communities.

We aim to develop an aeration-based strategy to enhance EPS production in activated sludge systems, building on the hypothesis that oxygen perturbation alters microbial metabolism, increases endogenous ROS generation, intensifies oxidative stress, and ultimately promotes EPS biosynthesis. However, the establishment of this strategy requires overcoming two key challenges: (i) understanding the impact of oxygen-driven oxidative stress on EPS biosynthesis in mixed microbial communities, and (ii) elucidating the mechanism of endogenous ROS generation under oxygen perturbations. Regarding the first challenge, it is known that activated sludge systems host diverse EPS-producing populations, primarily from the phyla *Proteobacteria, Nitrospirae*, and *Bacteroidetes* (Fang et al., 2024). These microbial groups not only dominate the biosystem but also possess strong antioxidant defenses (Johnson and Hug, 2019). However, it is unclear whether their capacity for rapid adaptation to oxidative environments contributes to the elevated EPS production observed in activated sludge. Moreover, the biosynthetic pathways and their types of EPS products can differ substantially among microbial species (Dueholm et al., 2023; Schmid, 2018). Consequently, the specific microbial responders and their associated biosynthetic activities can shape the overall EPS component profile, including the relative proportions of proteins, polysaccharides, and lipids, thereby influencing sludge characteristics such as floc structure and settleability, and the potential value of EPS for downstream recovery (Li et al., 2021).

As for the second challenge, the underlying mechanisms of endogenous ROS formation remain insufficiently understood. In microorganisms, ROS generation involves complex processes that extend beyond simple electron leakage from the respiratory chain (Seaver and Imlay, 2004). Seminal studies have proposed that the major contributors to ROS formation are flavoproteins that undergo unintended autoxidation (Ravindra Kumar and Imlay, 2013). These flavoproteins are distributed across multiple metabolic pathways, including but not limited to energy production, carbohydrate turnover, and amino acid metabolism. Representative enzymes include glutamate synthase, glutathione reductase, and lipoamide dehydrogenase (Imlay, 2013; Korshunov and Imlay, 2024). Other enzymes, such as proline dehydrogenase, which participates in the proline oxidation pathway, have been identified as a significant contributor to ROS generation (Zhang et al., 2015). These findings support the notion that variations in intracellular ROS levels are primarily linked to changes in core metabolic activity. While our previous work confirmed that amino acid metabolism is enhanced under oxygen perturbation (Zhou et al., 2025), it remains unclear how such metabolic shifts affect ROS production—particularly in terms of the generation enzymes, metabolic pathways, and taxonomic contributors involved.

To address these knowledge gaps, we examined the effects of three aeration regimes: continuous aeration (CA), continuous perturbation (CP), and intermittent perturbation (IP), each introducing a distinct oxygen exposure pattern. CA served as a stable control with constant oxygen supply, simulating conventional wastewater aeration. CP introduced high-frequency oxygen fluctuations under sustained aerobic conditions to intensify oxidative stress without inducing anoxia. Extending the CP regime, IP incorporated intermittent anoxic phases into the perturbation pattern to alleviate oxidative stress, thus enabling evaluation of whether oxidative stress was critical for enhancing EPS biosynthesis. Furthermore, we employed microbiome, metaproteome, and metabolome analyses to identify the microbial contributors and pathways involved in ROS generation and EPS production under these conditions. By elucidating the mechanisms through which oxygen perturbation induces oxidative stress and triggers microbial EPS biosynthesis as a response, this study provides a foundation for developing an aeration-driven strategy to enhance EPS production.

## 2. Materials and methods

### 2.1 Experimental setup and operation

Activated sludge used in this study was sourced from the Māngere Wastewater Treatment Plant in Auckland, New Zealand. Experiments were conducted in three identical cylindrical acrylic bioreactors (1□L working volume), operated in parallel at 20□°C under distinct aeration regimes. The same batch of activated sludge was used across all reactors, with three independent runs serving as biological triplicates. Each bioreactor was filled with 2.75□g/L mixed liquor suspended solids (MLSS), 3.84□g/L sodium bicarbonate (NaHCO□) as the inorganic carbon source, and 1□mL of trace element solution (composition detailed in Table S1). Over a 48-hour period, each system received a continuous and evenly distributed supply of 50□mL concentrated artificial wastewater (Table S2) via syringe pump, with magnetic stirring ensuring homogeneous mixing. MLSS, mixed liquor volatile suspended solids (MLVSS), sludge volume (SV) and sludge volume index (SVI) were analyzed according to the standard methods (Baird et al., 2017).

The three tested aeration conditions included: CA, maintaining dissolved oxygen (DO) at a stable level of 1.8– 2.2□mg/L (Figure S1); CP, characterized by DO oscillations between 0.1 and 2.0□mg/L in ~3-minute cycles (Figure S2); and IP, alternating between aerobic (DO up to 2.0□mg/L) and anoxic (DO near 0□mg/L) phases in 3-minute intervals (aerobic:anoxic = 3□min:3□min) (Figure S3).

### 2.2 EPS extraction and organic component analysis

EPS were extracted from activated sludge samples using a modified alkaline extraction protocol adapted from previous studies (Felz et al., 2019). Briefly, lyophilized sludge samples were suspended in 0.5% (w/v) sodium carbonate solution, with the total volume adjusted to ensure homogeneous mixing. The suspension was heated at 80□°C for 30□minutes under constant stirring to facilitate EPS release. After extraction, the mixture was centrifuged (4000 × g, 4□°C, 20□min) to remove solid residues. The supernatant was acidified to pH 2.2 using 1□M HCl to precipitate the EPS fraction, which was collected by centrifugation. The resulting EPS pellets were re-dissolved in 1□M NaOH (pH ~8.5) and subsequently subjected to desalting treatment to eliminate residual small molecules and salts. Desalted EPS samples were used directly for analysis.

Protein concentration was determined using the RC DC™ Protein Assay Kit I (Bio-Rad, USA), with bovine γ-globulin as the standard. Lipid content was analyzed via Soxhlet method (Abdulhussein Alsaedi et al., 2022), while carbohydrate content was measured using the phenol–sulfuric acid method with glucose as the calibration standard (Sheng et al., 2010).

### 2.3 ROS levels and antioxidant enzyme activities

To evaluate the oxidative stress responses under varying oxygen perturbation conditions, intracellular ROS levels were quantified at selected time intervals (0–8 h, 24–36 h, and at 48 h) using commercial fluorescence-based assays. The target ROS species—superoxide (O_2_•^-^), H_2_O_2_, and hydroxyl radical (•OH)—were detected using CellROX™ Green (Invitrogen, USA), OxiVision™ Green (AAT Bioquest, USA), and HPF Reagent (Invitrogen, USA), respectively. Reactions were carried out in 96-well microplates, with each well containing 50□μL of the working dye solution and 50□μL of the sample. For H_2_O_2_ detection, 10□μL of 0.3% sodium azide was added to suppress peroxidase interference. Following a 60-minute incubation in the dark at 20□°C, fluorescence was measured at an excitation/emission wavelength of 490/525□nm.

The activities of antioxidant enzymes were assessed to investigate cellular responses to ROS. Sludge samples (10□mL) were centrifuged at 5000 × g for 15 minutes at 4□°C, and the resulting pellets were washed twice with 50□mM Tris buffer. Cell disruption was achieved by ultrasonic treatment (40% amplitude, 10 seconds on/off cycles for 10 minutes) on ice using a QSONICA sonicator (USA). The extracted enzymes were subsequently analyzed for superoxide dismutase (SOD) and catalase (CAT) activities using commercially available assay kits (Abcam, UK), following the manufacturer’s instructions.

### 2.4 Metagenomic analysis

Metagenomic analysis with biological triplicates was conducted to identify key microbial gene contributors and to facilitate metaproteomic library development. Activated sludge samples collected at the 48-hour time point under aeration conditions were used for genomic DNA extraction, which was carried out using the DNeasy PowerSoil Kit (Qiagen, Germany) following the manufacturer’s protocol. High-throughput sequencing was performed on an Illumina HiSeq platform. Detailed information on sample preparation, sequencing procedures, and downstream bioinformatic processing is provided in Supplementary Materials (Section S1.3).

### 2.5 Metaproteomic analysis

Metaproteomic analysis was performed in biological triplicate to quantify protein expression and to identify key microbial enzymes contributors. Activated sludge samples were collected at the 48-hour time point under aerated conditions for protein extraction and downstream analysis. Proteins were isolated, purified, enzymatically digested, and subjected to nano LC-MS/MS using a TripleTOF 6600 mass spectrometer. Protein identification was carried out based on a custom database generated from the corresponding metagenomic data. Detailed protocols for protein extraction, sample preparation, and data processing are provided in Supplementary Materials (Section S1.4).

### 2.6 Metabolomic analysis

Intracellular metabolite profiling of activated sludge was carried out using gas chromatography–mass spectrometry (GC-MS) to investigate metabolic shifts under different experimental conditions. The analysis incorporated both biological and technical replicates to ensure data reliability. Metabolites were extracted and derivatized using the methyl chloroformate method. Comprehensive details of the extraction protocol, derivatization steps, GC-MS analysis, and data processing are provided in Supplementary Materials (Section S1.5).

## 3. Results

### 3.1. Modulating EPS accumulation and composition through continuous oxygen perturbation

#### 3.1.1. Increased EPS yield without compromising treatment performance

Wastewater treatment performance remained comparable under CA and CP, showing that oxygen perturbation did not compromise organic matter removal or ammonia oxidation. The carbon oxidation rates were nearly identical under CA (75.4 ± 1.4 mg COD/L/hr) and CP (75.5 ± 1.7 mg COD/L/hr) conditions (Table 1). Similarly, nitrification rates remained stable under both CA and CP conditions, measured at 7.8 ± 0.3 and 7.8 ± 0.2 mg NH_4_^+^-N/L/hr, respectively. In addition, denitrification rates were very low under both CA (0.01 ± 0.01 mg NO_x_^-^-N/L/hr) and CP (0.73 ± 0.08 mg NO_x_^-^-N/L/hr), suggesting that the high-frequency oxygen switching in CP did not significantly restrict microbial access to oxygen. However, under IP, the system exhibited distinct shifts in nitrogen transformation behavior. Nitrification rates dropped to 5.2 ± 0.4 mg NH_4_^+^-N/L/hr, while denitrification rates increased significantly to 3.3 ± 0.1 mg NO_X_^-^-N/L/hr. These shifts indicate a transition toward oxygen-limited conditions under IP, as evidenced by decreased nitrification and increased denitrification rates.

**Table 1.**
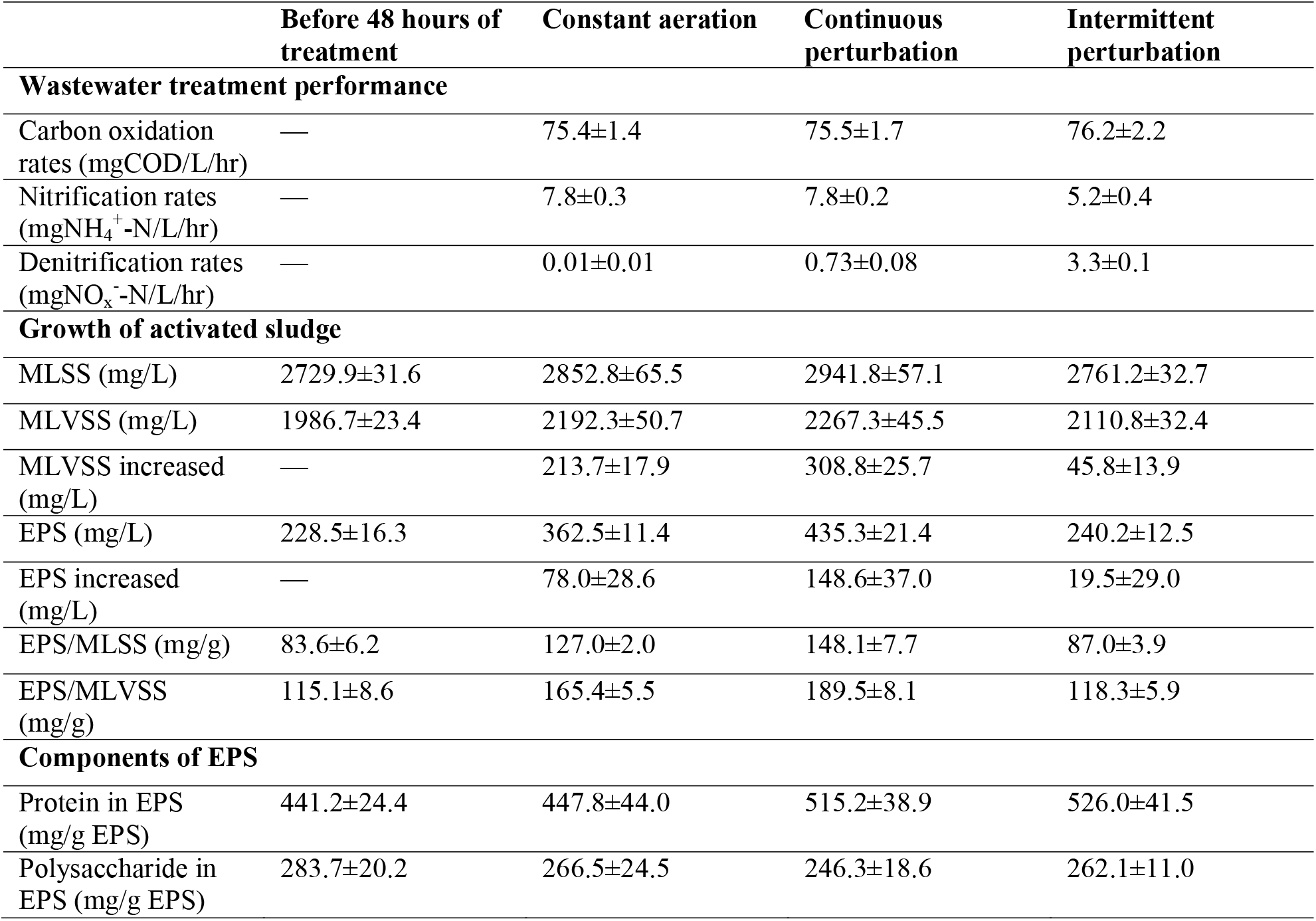
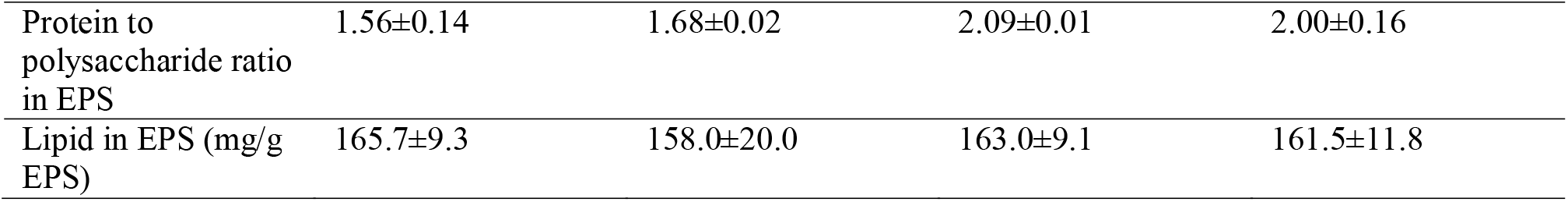
Wastewater treatment performance, activated sludge growth, and EPS composition before and after 48 hours of exposure to different aeration conditions.

|  | Before 48 hours of treatment | Constant aeration | Continuous perturbation | Intermittent perturbation |
| --- | --- | --- | --- | --- |
| **Wastewater treatment performance** |  |  |  |  |
| Carbon oxidation rates (mgCOD/L/hr) | — | $75.4 \pm 1.4$ | $75.5 \pm 1.7$ | $76.2 \pm 2.2$ |
| Nitrification rates (mgNH<sub>4</sub><sup>+</sup>-N/L/hr) | — | $7.8 \pm 0.3$ | $7.8 \pm 0.2$ | $5.2 \pm 0.4$ |
| Denitrification rates (mgNO<sub>x</sub><sup>-</sup>-N/L/hr) | — | $0.01 \pm 0.01$ | $0.73 \pm 0.08$ | $3.3 \pm 0.1$ |
| **Growth of activated sludge** |  |  |  |  |
| MLSS (mg/L) | $2729.9 \pm 31.6$ | $2852.8 \pm 65.5$ | $2941.8 \pm 57.1$ | $2761.2 \pm 32.7$ |
| MLVSS (mg/L) | $1986.7 \pm 23.4$ | $2192.3 \pm 50.7$ | $2267.3 \pm 45.5$ | $2110.8 \pm 32.4$ |
| MLVSS increased (mg/L) | — | $213.7 \pm 17.9$ | $308.8 \pm 25.7$ | $45.8 \pm 13.9$ |
| EPS (mg/L) | $228.5 \pm 16.3$ | $362.5 \pm 11.4$ | $435.3 \pm 21.4$ | $240.2 \pm 12.5$ |
| EPS increased (mg/L) | — | $78.0 \pm 28.6$ | $148.6 \pm 37.0$ | $19.5 \pm 29.0$ |
| EPS/MLSS (mg/g) | $83.6 \pm 6.2$ | $127.0 \pm 2.0$ | $148.1 \pm 7.7$ | $87.0 \pm 3.9$ |
| EPS/MLVSS (mg/g) | $115.1 \pm 8.6$ | $165.4 \pm 5.5$ | $189.5 \pm 8.1$ | $118.3 \pm 5.9$ |
| **Components of EPS** |  |  |  |  |
| Protein in EPS (mg/g EPS) | $441.2 \pm 24.4$ | $447.8 \pm 44.0$ | $515.2 \pm 38.9$ | $526.0 \pm 41.5$ |
| Polysaccharide in EPS (mg/g EPS) | $283.7 \pm 20.2$ | $266.5 \pm 24.5$ | $246.3 \pm 18.6$ | $262.1 \pm 11.0$ |
| Protein to polysaccharide ratio in EPS | 1.56±0.14 | 1.68±0.02 | 2.09±0.01 | 2.00±0.16 |
| Lipid in EPS (mg/g EPS) | 165.7±9.3 | 158.0±20.0 | 163.0±9.1 | 161.5±11.8 |

The activated sludge responded differently in terms of biomass growth and EPS accumulation under the various aeration regimes. Elevated MLSS and MLVSS concentrations under both CA and CP conditions reflect enhanced microbial proliferation and biomass accumulation. The observed biomass increase in activated sludge was primarily attributed to the accumulation of organic matter, with EPS contributing a significant portion. After 48 hours of operation, MLVSS under CP increased from 1986.7 ± 23.4 mg/L (initial) to 2267.3 ± 45.5 mg/L, corresponding to a biomass gain of 308.8 ± 25.7 mg/L (Table 1). This increase was 45% greater than that observed under CA, which can be partially attributed to the increased contribution of EPS. Under CP conditions, EPS increased by 148.6 ± 37.0 mg/L, approximately 90% higher than the increase observed under CA (78.0 ± 28.6 mg/L). In contrast, IP conditions resulted in the lowest EPS accumulation, with total EPS content reaching 240.2 ± 12.5 mg/L and a net increase of only 19.5 ± 29.0 mg/L. This limited increase implies that the oxygen fluctuations and extended anoxic phases under IP were insufficient to promote significant EPS or organic biomass accumulation. Ultimately, the proportion of EPS relative to MLVSS in the treated activated sludge was highest under CP conditions, with an EPS/MLVSS ratio of 189.5 ± 8.1 mg/g, compared to 165.4 ± 5.5 mg/g for CA and 118.3 ± 5.9 mg/g for IP (Figure 1a).

**Figure 1.**
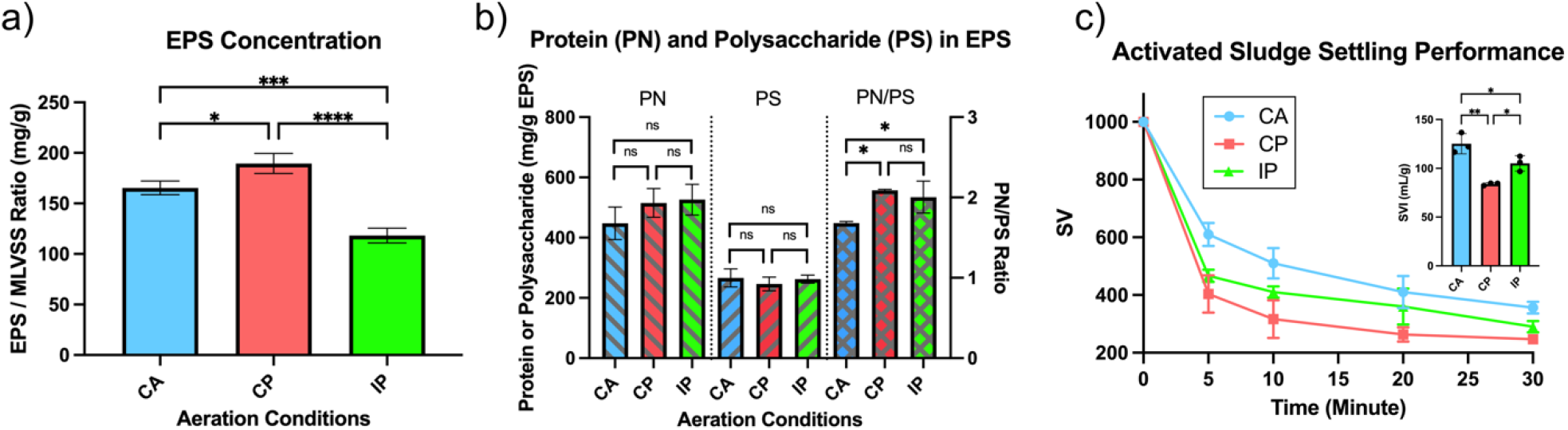
Enhanced EPS yield and PN/PS ratio improve sludge settleability under oxygen perturbation conditions. (a) EPS/MLVSS ratio under constant aeration (CA), continuous perturbation (CP), and intermittent perturbation (IP) after 48 h; (b) EPS composition and protein-to-polysaccharide ratio; (c) Sludge volume (SV) and sludge volume index (SVI) across conditions. *Error bars represent SD (n = 3 biological replicates); significance tested by one-way ANOVA*.

#### 3.1.2. Enhanced EPS protein-to-polysaccharide ratio improves sludge settleability

The primary EPS components of interest for resource recovery are proteins, polysaccharides, and lipids, which together represent the major recoverable organic fractions in activated sludge. Targeted compositional analysis further confirmed that proteins were the predominant component of EPS, accounting for approximately half of the total mass. Under CA, CP, and IP conditions, the protein content in EPS was 447.8 ± 44.0, 515.2 ± 38.9, and 526.0 ± 41.5 mg/g EPS, respectively (Table 1, Figure 1b). Polysaccharides were the second most abundant EPS component, accounting for roughly one-quarter of the total, with levels of 266.5 ± 24.5, 246.3 ± 18.6, and 262.1 ± 11.0 mg/g EPS under CA, CP, and IP, respectively. Lipids were less abundant than proteins and polysaccharides, with concentrations of 158.0 ± 20.0, 163.0 ± 9.1, and 161.5 ± 11.8 mg/g EPS under CA, CP, and IP conditions, respectively.

Although the individual contents of proteins, polysaccharides, and lipids in EPS did not differ significantly among the three aeration conditions based on statistical analysis, the protein-to-polysaccharide (PN/PS) ratio emerged as a distinct characteristic, showing markedly higher values under CP and IP conditions compared to CA (Table 1, Figure 1b). The elevated PN/PS ratio under oxygen perturbation conditions reflects that protein production outpaced that of polysaccharides, contributing to improved sludge flocculation. This finding is consistent with the observed differences in sludge settling performance (Figure 1c). Sludge treated under CP and IP settled more rapidly than under CA, with SVI values of 125 ± 8 mL/g for CA and a significantly lower SVI of 83 ± 1 mL/g under CP, showing better settleability.

### 3.2. Oxidative stress triggers antioxidant response and EPS formation under continuous oxygen perturbation

#### 3.2.1. Elevated H_2_O_2_ and catalase activity indicate intensified oxidative stress

This section reanalyzes previously reported data to explore the relationship between oxidative stress and EPS formation under different aeration conditions. During the 48-hour bioreactor operation, distinct differences in H_2_O_2_ concentrations were observed under the three aeration conditions (Figure 2a). In contrast, concentrations of other ROS, including O_2_•^-^ and •OH, did not differ significantly (Figure S4, S5), indicating that the accumulation of H_2_O_2_ was the primary ROS behind the oxidative stress. CP resulted in the highest intracellular H_2_O_2_ levels, followed by CA, whereas IP consistently maintained the lowest concentrations throughout the experiment (*P*<0.05, repeated-measures ANOVA with Tukey’s post hoc test). Under both CA and CP conditions, H_2_O_2_ levels initially increased and then stabilized, reaching steady-state ranges of 21–45 μM and 32–56 μM, respectively. In contrast, concentrations under IP remained consistently below 20 μM. These results indicate that continuous oxygen perturbation effectively elevated intracellular H_2_O_2_ concentrations and confirmed stronger oxidative stress under CP conditions.

**Figure 2.**
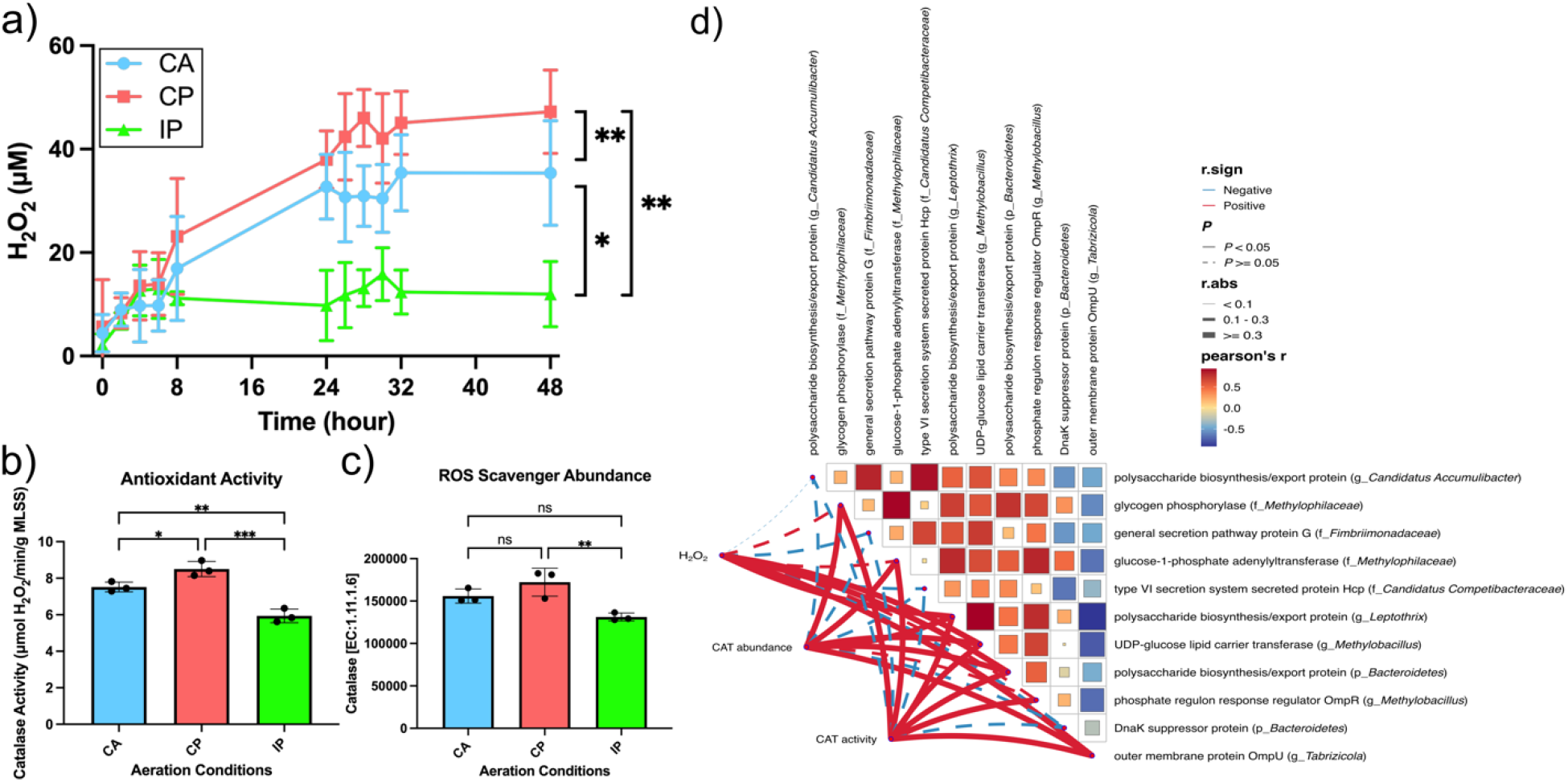
H_2_O_2_ accumulation and catalase induction indicate oxidative stress under continuous oxygen perturbation. (a) Time-course of intracellular H_2_O_2_ concentrations under different aeration modes; (b) Catalase (CAT) activity measured by H_2_O_2_ degradation rate; (c) Relative protein abundance of CAT from metaproteomic data; (d) Pearson correlation network linking oxidative stress (H_2_O_2_, CAT abundance and activity) with biofilm-related proteins. *Error bars represent SD (n = 3 biological replicates); significance tested by repeated-measures ANOVA followed by Tukey’s multiple comparisons test for (a); significance tested by one-way ANOVA for (b) and (c)*.

To evaluate microbial antioxidant responses, we analyzed both the activity and abundance of two key antioxidant enzymes: SOD, which scavenges O_2_•^-^, and CAT, which decomposes H_2_O_2_. No significant variation in SOD activity or protein abundance was observed among the different aeration conditions (Figure S6, S7). In contrast, CAT activity and protein abundance were highest under CP conditions (Figure 2b and 2c), consistent with the elevate H_2_O_2_ levels observed in this group. A moderate upregulation of CAT was observed under CA, whereas IP maintained the lowest CAT levels with minimal variation throughout the experimental period. Correspondingly, CAT-mediated H_2_O_2_ scavenging capacity reached 8.5 ± 0.3 μM/min/g under CP, significantly higher than that under CA (7.5 ± 0.2 μM/min/g) and IP (5.9 ± 0.3 μM/min/g).

#### 3.2.2. H_2_O_2_-linked protein associations suggest potential connections to EPS-related functions

To investigate the potential link between oxidative stress and EPS formation, a Pearson correlation analysis was performed between three oxidative indicators (H_2_O_2_, CAT abundance, and CAT activity) and the abundance of biofilm-related proteins. These proteins, originating from diverse microbial taxa, were prefiltered based o significant variation across aeration conditions (ANOVA, p < 0.05). As shown in Figure 2d, the analysis identified significant positive correlations (r > 0.3, p < 0.05), with CAT activity exhibiting the highest number of associations (7 proteins), followed by CAT abundance (6 proteins), and H_2_O_2_ concentration (4 proteins). Proteins positively associated with H_2_O_2_ included two polysaccharide biosynthesis/export proteins from *Bacteroidetes* and *Leptothrix*, a UDP-glucose lipid carrier transferase from *Methylobacillus*, and the outer membrane protein OmpU from *Tabrizicola*. In addition to these, proteins significantly correlated with CAT abundance included glycogen phosphorylase from *Methylophilaceae* and glucose-1-phosphate adenylyltransferase from *Methylobacillus*. All these proteins also showed significant correlations with CAT activity, which further exhibited a unique positive association with the phosphate regulon response regulator OmpR from *Methylobacillus*.

### 3.3. Flavoenzyme-associated pathways as sources of ROS under continuous oxygen perturbation

To trace the metabolic origins of endogenous ROS, we identified flavoenzymes in the metaproteomic dataset. Among the 88 identified flavoenzymes, we focused specifically on those classified under the “metabolism” category at KEGG level 1 (Figure S8). A total of 60 such metabolism-related flavoenzymes were identified, which were distributed across 10 KEGG level 2 pathways. Several enzymes were involved in multiple metabolic routes. Among these, energy metabolism contained the highest number of flavoenzymes (21), followed by carbohydrate metabolism (18) and amino acid metabolism (15) (Figure 3). This distribution pattern suggests that ROS production primarily originates from core microbial metabolic activities, particularly those related to central carbon and nitroge metabolism.

**Figure 3.**
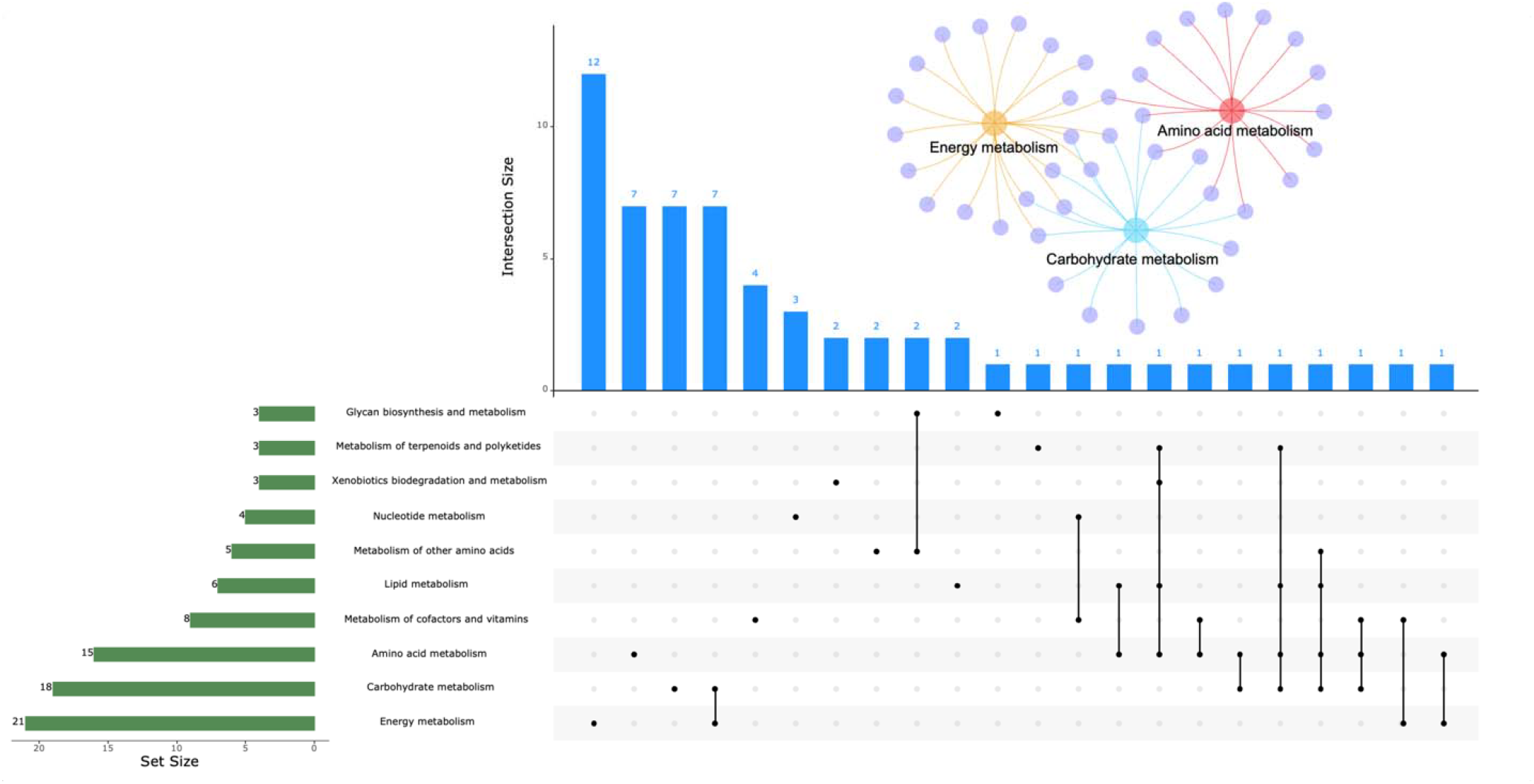
Distribution of identified flavoenzymes across central microbial metabolic pathways. KEGG level 2 pathway classification of 60 flavoenzymes involved in metabolic processes with ROS-generation potential, identified from metaproteomic data. Energy, carbohydrate, and amino acid metabolism dominate flavoenzyme localization, implicating them as major hubs for redox overflow.

To further identify the metabolic drivers of elevated ROS accumulation under CP conditions, we analyzed all 6 flavoenzymes and selected a subset based on significant changes in the concentrations of their electron-donating substrates (Supplementary Table S3, Excel format). Electron-donating substrates provide reducing equivalents to flavoenzymes, which can lead to unintended electron leakage to molecular oxygen, thereby generating ROS as byproducts. Although no significant differences in protein abundance for these flavoenzymes were observed between CA and CP conditions, the concentrations of their electron-donating substrates consistently increased under CP, including glutamate, glutathione, glycine, proline, isoleucine, valine, and leucine. These metabolites are associated with multiple ROS-generating flavoenzymes identified under CP conditions. Among these enzymes, glutamate synthase (NADPH), glutathione reductase, and dihydrolipoamide dehydrogenase were consistently detected across all biological replicates (Figure 4), whereas glutamate synthase (ferredoxin), proline dehydrogenase, and leucine dehydrogenase were identified in a subset of samples (Figure S9).

**Figure 4.**
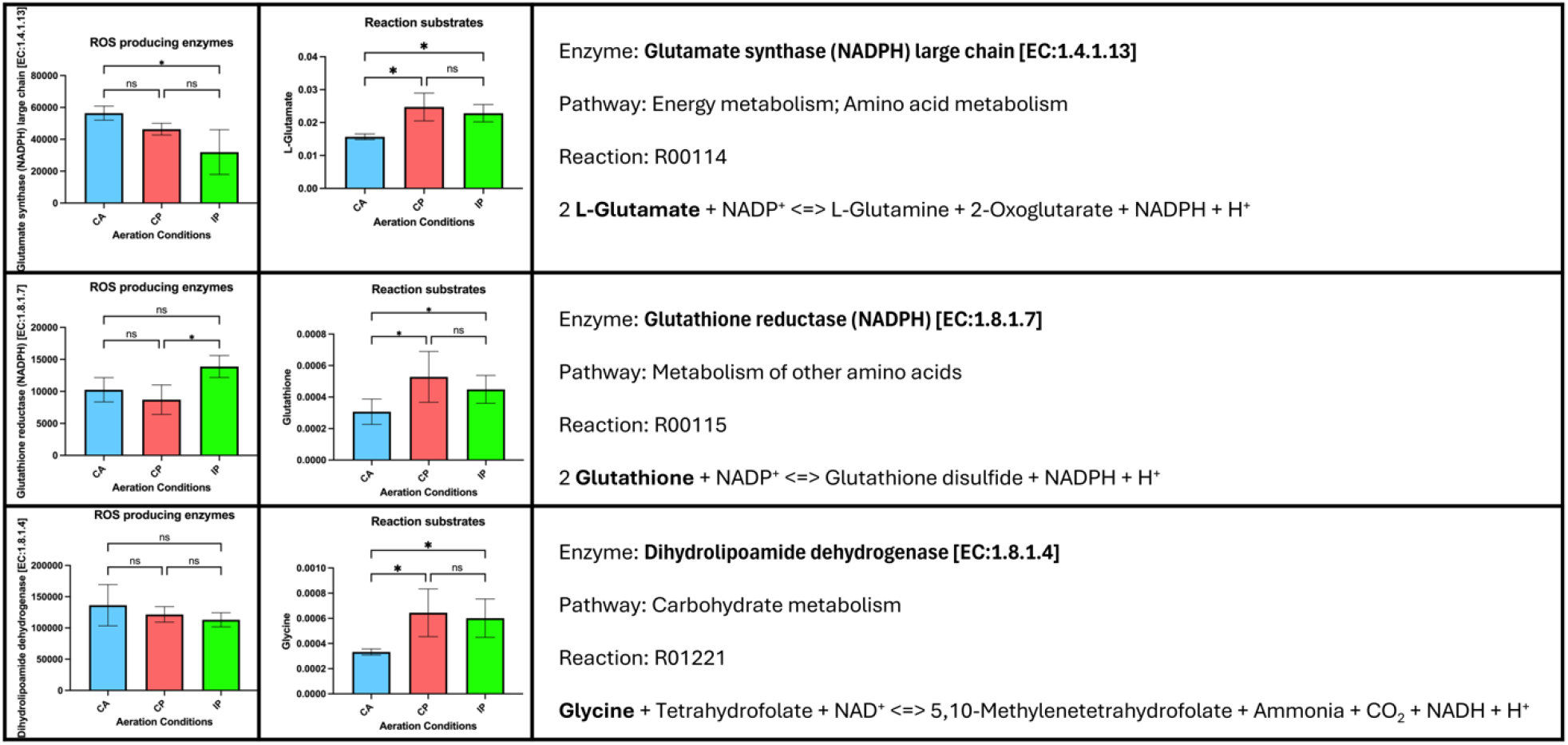
Candidate flavoenzymes contributing to ROS production via elevated metabolite accumulation under CP. Relative protein abundance of three ROS-related flavoenzymes and corresponding electron-donating substrate concentrations under CA, CP, and IP. Highlighted enzymes include glutamate synthase (NADPH), glutathione reductase, and dihydrolipoamide dehydrogenase. *Error bars represent standard deviation; n = biological replicates*.

Under IP conditions, these electron-donating substrates also showed elevated concentrations, similar to those under CP (Figure 4, Figure S9), implying that oxygen perturbation in general may increase the potential for electro leakage in flavoenzyme-associated pathways. However, ROS levels were significantly lower under IP than CP. This discrepancy suggests that while metabolic activity may promote conditions favorable to ROS formation, the actual generation of ROS requires sufficient oxygen availability to serve as an unintended electron acceptor.

### 3.4. Microbial contributors to endogenous ROS production

#### 3.4.1. Taxonomic diversity in ROS gene contributors

To understand the ecological basis of intracellular ROS generation, metagenomic data were used to identify microbial contributors harboring genes encoding six ROS-generating enzymes: glutamate synthase (NADPH) (K00265), glutathione reductase (K00383), dihydrolipoamide dehydrogenase (K00382), glutamate synthase (ferredoxin) (K00284), proline dehydrogenase (K00318), and leucine dehydrogenase (K00263). Taxonomic distributions at the family level under CA, CP, and IP conditions were characterized and compared (Figures S10 S27). Focusing only on taxonomically resolved families, we applied a cumulative contribution threshold of 95% to define major microbial contributors under CP. Functional redundancy varied among the six genes: K00265 an K00382 exhibited high redundancy, requiring 13 and 18 families, respectively, to reach the threshold. In contrast, K00383 and K00263 showed more specialized patterns, with only 4 and 5 families contributing, respectively. Intermediate diversity was observed in K00284 (7 families) and K00318 (10 families) (Figure 5).

**Figure 5.**
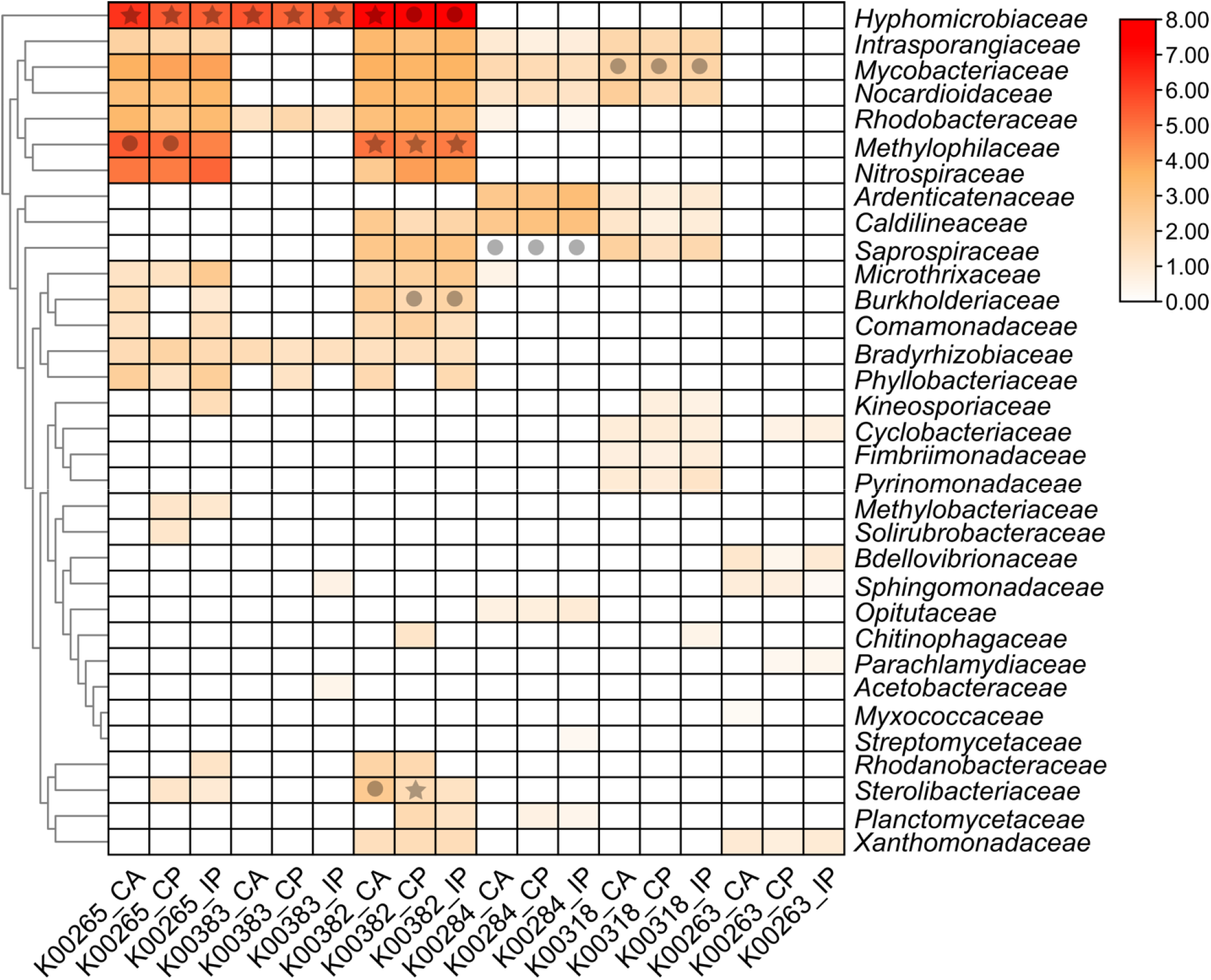
Family-level heatmap of microbial contributors to ROS-generation genes and enzymes expressed. Heatmap values represent the relative contribution of taxonomically identified families accounting for the top 95% cumulative contribution of each gene (K00265, K00383, K00382, K00284, K00318, and K00263) under three aeration regimes: CA, CP, and IP. *Markers denote protein detection consistency across biological replicates. ⍰ indicates families with consistent expression (detected in all three replicates);* • *indicates partial detection (1– replicates)*

Comparison across ROS-generating genes revealed a group of microbial families that consistently contributed to multiple genes. Among them, *Mycobacteriaceae, Nocardioidaceae*, and *Intrasporangiaceae* were involved in four or more ROS-related genes, underscoring their central ecological roles. In parallel, families such as *Hyphomicrobiaceae, Methylophilaceae, Nitrospiraceae*, and *Rhodobacteraceae* were predominantly associated with broad-function genes like K00265 and K00382, while also serving as key contributors within their respective metabolic niches. Additionally, *Hyphomicrobiaceae* accounted for 84% of the gene abundance of K00383, illustrating its dual role in both broad-spectrum and pathway-specific ROS-associated functions. In contrast, the microbial contributions to K00284, K00318, and K00263 were relatively lower, likely due to the lack of support from abundant families such as *Hyphomicrobiaceae* and *Methylophilaceae*.

Notably, among the major contributors to ROS-generating genes, several families corresponded to microbial genera previously identified as upregulating EPS biosynthesis proteins under oxidative stress. Specifically, the genera *Methylobacillus* (family *Methylophilaceae*), *Tabrizicola* (family *Rhodobacteraceae*), and *Leptothrix* (family *Comamonadaceae*) were associated with contributions to key ROS-generating genes. *Methylobacillus* contributed to K00265 and K00382, while *Tabrizicola* was linked to K00382. Although *Leptothrix* itself was not directly detected at the genus level in ROS contributors, its parent family, *Comamonadaceae*, contributed to both K00265 and K00382. These overlaps suggest that oxidative stress responses linked to ROS generation may stimulate EPS biosynthesis at the genus level, although it remains unclear whether the same finer-scale microbial taxa are simultaneously responsible for both processes.

#### 3.4.2. Cross-validation of ROS-producer by gene and protein evidence

Metaproteomics provided valuable confirmation of enzyme expression and a complementary perspective on microbial origins. The expression of six ROS-generating flavoenzymes was mainly affiliated with *Proteobacteria*— particularly *Alphaproteobacteria* and *Betaproteobacteria*—along with members of *Bacteroidetes* and *Actinobacteria* (Table S4). Specifically, glutamate synthase (NADPH), glutathione reductase, and dihydrolipoamide dehydrogenase were predominantly derived from *Proteobacteria*, whereas glutamate synthase (ferredoxin) and leucine dehydrogenase were mainly associated with *Bacteroidetes*, and proline dehydrogenase with *Actinobacteria*.

Within the framework of taxonomic contributors identified via gene analysis, *Hyphomicrobiaceae* expressed glutamate synthase (NADPH), glutathione reductase, and dihydrolipoamide dehydrogenase, while *Methylophilaceae* expressed both glutamate synthase (NADPH) and dihydrolipoamide dehydrogenase (Figure 5). Within *Methylophilaceae*, Methylobacillus, previously identified as upregulating EPS biosynthesis proteins in response to oxidative stress, was confirmed to express glutamate synthase (NADPH) at levels sufficient for detection and taxonomic resolution (Table S4). For dihydrolipoamide dehydrogenase, additional contributors were observed in *Burkholderiaceae* and *Sterolibacteriaceae*, with expression patterns showing sensitivity to aeration conditions. Although *Mycobacteriaceae* contributed gene sequences for four flavoenzymes, only proline dehydrogenase was detected in the protein profile. An exception was observed for *Saprospiraceae*, which expressed glutamate synthase (ferredoxin) despite no confirmed gene contribution. This mismatch between gene potential and protein detection likely reflects the current limitations in resolving microbial activity at fine taxonomic scales. Nevertheless, the combined data provide a functional snapshot of potential ROS-producing microbial taxa under oxygen perturbation.

## 4. Discussion

### 4.1. A mechanistic framework connecting ROS and EPS under oxygen perturbation

This study establishes a mechanistic connection linking oxygen perturbation, intracellular oxidative stress, and enhanced extracellular polymeric substances production in activated sludge. While it is widely recognized that environmental stressors can trigger microorganisms to secrete more EPS as a protective mechanism (Shi et al., 2017), demonstrating a clear cause-effect sequence in complex microbial communities has remained challenging. Here, by applying a controlled redox fluctuation regime, we observed a reproducible oxidative stress response, evidenced by H_2_O_2_ accumulation and the activation of antioxidant enzymes, alongside enhanced extracellular polymeric substances biosynthesis.

The positive correlations among H_2_O_2_, CAT abundance/activity, and multiple EPS-related proteins support the notion that oxidative stress induces a coordinated upregulation of exopolysaccharide biosynthesis across diverse microbial taxa. The enzymes identified span multiple stages of carbon metabolism linked to EPS production. For example, glycogen phosphorylase and glucose-1-phosphate adenylyltransferase, both detected in *Methylophilaceae*, participate in the mobilization and conversion of glucose-1-phosphate, a central metabolic intermediate that connects glycogen turnover with the synthesis of UDP-glucose precursors for EPS biosynthesis and contributes to glycogen metabolic balance (Mao et al., 2006). Meanwhile, UDP-glucose lipid carrier transferases from *Methylobacillus* and polysaccharide biosynthesis/export proteins from *Bacteroidetes* and *Leptothrix* represent core elements of the Wzx/Wzy-dependent pathway, which assembles and exports branched heteropolysaccharides via lipid-linked intermediates (Dong et al., 2006; Schmid, 2018). The observed correlation between OmpR and CAT activity further supports the role of H_2_O_2_ as a regulatory signal that modulates both redox defense and exopolysaccharide production through transcriptional cascades. OmpR is a response regulator activated via phosphorylation by EnvZ under environmental stimuli such as osmotic or oxidative stress (Cai and Inouye, 2002, Seo et al., 2017). Upon activation, OmpR binds DNA as a tandem dimer and further facilitates transcriptional regulation of stress-responsive genes, including those involved in porin expression and potentially EPS biosynthesis under oxidative conditions (Harrison-McMonagle et al., 1999; Prigent-Combaret et al., 2001).

To our knowledge, this is the first study to comprehensively map this chain of events under oxygen perturbation in an ecologically realistic system (activated sludge). While previous reports have acknowledged the presence of reactive oxygen species during redox fluctuations and their potential role in pollutant degradation, their connection to biopolymer production remains unclear (Bains et al., 2019; Liu et al., 2024). Other studies have observed elevated EPS production under pure oxygen aeration; however, the involvement of oxidative stress as a mechanistic driver has remained largely speculative (Hu et al., 2019). Here, by measuring ROS and linking them to specific metabolic sources (flavoenzymes) and EPS outcomes, we bridge the gap between these phenomena. We support the idea that oxidative stress is not merely a byproduct of oxygen perturbation; instead, it functions as the central driver that redirects metabolic resources towards EPS synthesis. This represents a paradigm shift from viewing oxidative stress as purely detrimental to recognizing it as a controllable trigger for beneficial outcomes. This concept of “oxidative eustress” has broad implications, as it highlights the potential for operational strategies in wastewater bioprocesses can target microbial stress responses to enhance specific functions (Sies, 2021).

### 4.2. Flavoenzyme-driven ROS generation and microbial response to oxygen perturbation

CP primarily promoted increased ROS generation via non-respiratory flavoenzymes rather than through classical electron transport chain (ETC) leakage. Traditionally, ROS are viewed as byproducts of respiration arising from electron leakage at ETC complexes I and III (Messner and Imlay, 1999). However, our metaproteomic data showed no significant increase in respiratory enzyme expression under CP. Instead, metabolomic analysis reflected that flavoenzymes involved in amino acid metabolism such as glutamate synthase, glutathione reductase, dihydrolipoamide dehydrogenase, proline dehydrogenase, and leucine dehydrogenase contributed substantially to the elevated ROS levels observed under CP conditions, consistent with the latest understanding of ROS-generating enzymes (Fleminger and Dayan, 2021; Imlay, 2013; Korge et al., 2015; Korshunov and Imlay, 2024; Yang et al., 2023; Zhang et al., 2015). The accumulation of their electron-donating substrates including glycine, proline, glutamate, and branched-chain amino acids regarded as enhanced substrate utilization, leading to redox overflow and subsequent ROS formation. Among the measured ROS species, H_2_O_2_ showed the most significant increase under CP, which correlated closely with increased catalase activity, highlighting H_2_O_2_ as the dominant oxidative stress indicator. In contrast, O_2_•^-^ and •OH remained unchanged, supporting the view that elevated H_2_O_2_ primarily originated from the direct two-electron reduction of oxygen by flavoenzymes under CP conditions (Ravindra Kumar and Imlay, 2013; Seaver and Imlay, 2004).

Metagenomic analyses detected a wide range of microbial taxa harboring genes encoding ROS-generating flavoenzymes. Generalist enzymes such as glutamate synthase and dihydrolipoamide dehydrogenase exhibited high functional redundancy across multiple microbial families (Chen et al., 2022). In contrast, specialized enzymes like glutathione reductase and leucine dehydrogenase showed narrower taxonomic distributions, pointing to more constrained metabolic roles (Chen et al., 2021). Metaproteomic data further underscored *Hyphomicrobiaceae* and *Methylophilaceae* as dominant contributors to flavoenzyme expression, likely functioning as basal ROS producers due to their broad metabolic versatility. Notably, *Methylophilaceae*, which was identified as a key contributor to ROS generation, also exhibited increased expression of EPS biosynthesis-related enzymes under CP conditions. This dual functional role suggests that *Methylophilaceae* may directly couple redox overflow with EPS production, reinforcing the metabolic linkage between oxidative stress and biosynthetic responses triggered by oxygen perturbation (Deng et al., 2024).

Despite significant increases in electron-donating substrates under CP, the abundances of ROS-generating enzymes remained largely unchanged across aeration conditions. This confirms that enhanced ROS accumulation was not driven by changes in flavoenzyme expression. Although substrate levels were similarly elevated under both CP and IP conditions, only CP resulted in sustained ROS accumulation, indicating oxygen availability as a key determinant. Our previous study demonstrated that microbial communities primarily responded to oxygen perturbation by enhancing amino acid biosynthesis and redistributing metabolic fluxes (Zhou et al., 2025). In this study, these responses are considered key drivers of oxidative stress, rather than substantial changes in ROS-related community composition or flavoenzyme expression.

### 4.3. Limitations and future perspectives

In contrast to external or resource-intensive interventions, the CP strategy adopted in this study leverages an operational variable inherently present in wastewater treatment, inducing stress simply by adjusting the aeration cycle without requiring additional chemicals or substrates, offering greater environmental friendliness and controllability. Moreover, stable treatment performance and improved sludge settleability further support the feasibility of scaling this strategy to large systems. However, current experiments were conducted in bench-scale reactors under controlled conditions. Scaling up CP in full-scale WWTPs may present challenges, including aeration control complexity and sensor-based feedback requirements. Despite these issues, EPS extraction from CAS under CP reached 189.5□mg/g MLVSS, comparable to the lower range of values reported for EPS recovered from aerobic granular sludge (Li et al., 2021). It is conservatively estimated that implementing CP in a municipal WWTP with a capacity of 12,000□m^3^/day could generate an additional economic benefit of approximately $193,500 USD annually through enhanced EPS recovery (Supplementary Materials Section S2.4).

This study also provides mechanistic insights into how oxygen perturbation enhances EPS production by modulating intracellular redox dynamics and stimulating flavoenzyme-driven ROS formation. However, several limitations should be acknowledged. Although metaproteomic and metabolomic data allowed inference of ROS-generating pathways and microbial contributors, current omics resolutions may not fully capture low-abundance proteins or rare taxa potentially critical. Moreover, microbial communities vary in composition and metabolic capacity across WWTPs, and the key microbial responders involved in EPS biosynthesis under CP conditions may differ between treatment plants. Such differences could lead to variations in EPS product profiles, including the protein-to-polysaccharide ratio and the presence of specific structural components, necessitating system-specific evaluations.

Future work should focus on: (i) long-term studies evaluating the ecological stability and process robustness of CP under dynamic influent conditions and seasonal fluctuations; (ii) techno-economic assessments of EPS recovery at pilot and full scales to determine its cost-effectiveness and environmental benefits; and (iii) comparative analyses of microbial communities across different WWTPs to identify universal versus site-specific EPS-producing taxa and biosynthetic pathway preferences under CP strategy.

## 5. Conclusion

This study highlights that deliberate modulation of microbial oxidative stress via oxygen perturbation offers a promising route toward sustainable resource recovery in wastewater treatment systems. By shifting from reliance on external chemical inputs to exploiting inherent microbial metabolic adaptability, this approach presents a more sustainable and scalable alternative. The results emphasize the potential of ecological disturbance as an operational tool, broadening the scope for integrating microbial ecology into practical engineering solutions and providing a fresh perspective on advancing wastewater resource management strategies.

## Abbreviations

Abbreviation Full Form

EPS: Extracellular Polymeric Substances
CP: Continuous Perturbation
WWTPs: Wastewater Treatment Plants
CAS: Conventional Activated Sludge
ROS: Reactive Oxygen Species
CA: Continuous Aeration
IP: Intermittent Perturbation
MLSS: Mixed Liquor-Suspended Solid
MLVSS: Mixed Liquor Volatile Suspended Solids
SV: Sludge Volume
SVI: Sludge Volume Index
DO: Dissolved Oxygen
O_2_•^-^: Superoxide
•OH: Hydroxyl Radical
SOD: Superoxide Dismutase
CAT: Catalase
PN/PS: ratio Protein-to-Polysaccharide Ratio
ETC: Electron Transport Chain

## Corresponding Author

* Phone: +64 9 923 4512; fax: +64 9 373 7462.

## Acknowledgments

We thank Martin Middleditch, George Guo, Saras Green, and Alastair Harris for their assistance with metaproteomics and metabolomics analysis, Watercare Services Ltd for activated sludge culture, and acknowledge NeSI facilities and the University of Auckland’s Centre for eResearch for their support.

## Supporting Information

All the supplementary tables and figures are provided in Supporting Information.

## Reference

Abdulhussein Alsaedi, A., Sohrab Hossain, Md., Balakrishnan, V., Abdul Hakim Shaah, M., Mohd Zaini Makhtar, M., Ismail, N., Naushad, Mu., Bathula, C., 2022. Extraction and separation of lipids from municipal sewage sludge for biodiesel production: Kinetics and thermodynamics modeling. Fuel 325, 124946. 10.1016/j.fuel.2022.124946

Bahgat, N.T., Wilfert, P., Korving, L., van Loosdrecht, M., 2023. Integrated resource recovery from aerobic granular sludge plants. Water Research 234, 119819. 10.1016/j.watres.2023.119819

Bains, A., Perez-Garcia, O., Lear, G., Greenwood, D., Swift, S., Middleditch, M., Kolodziej, E.P., Singhal, N., 2019. Induction of microbial oxidative stress as a new strategy to enhance the enzymatic degradation of organic micropollutants in synthetic wastewater. Environ. Sci. Technol. 53, 9553–9563. 10.1021/acs.est.9b02219

Baird, R., Bridgewater, L., American Public Health Association, American Water Works Association, Water Environment Federation (Eds.), 2017. Standard methods for the examination of water and wastewater, 23rd edition. ed. American Public Health Association, Washington, D.C.

Cai, S.J., Inouye, M., 2002. EnvZ-OmpR interaction and osmoregulation in Escherichia coli. Journal of Biological Chemistry 277, 24155–24161. 10.1074/jbc.M110715200

Chen, Huaihai, Ma, K., Lu, C., Fu, Q., Qiu, Y., Zhao, J., Huang, Y., Yang, Y., Schadt, C.W., Chen, Hao, 2022. Functional redundancy in soil microbial community based on metagenomics across the globe. Front. Microbiol. 13. 10.3389/fmicb.2022.878978

Chen, Y.-J., Leung, P.M., Wood, J.L., Bay, S.K., Hugenholtz, P., Kessler, A.J., Shelley, G., Waite, D.W., Franks, A.E., Cook, P.L.M., Greening, C., 2021. Metabolic flexibility allows bacterial habitat generalists to become dominant in a frequently disturbed ecosystem. The ISME Journal 15, 2986–3004. 10.1038/s41396-021-00988-w

Deng, H., Li, Q., Li, M., Sun, L., Li, B., Wang, Y., Wu, Q.L., Zeng, J., 2024. Epiphytic microorganisms of submerged macrophytes effectively contribute to nitrogen removal. Environmental Research 242, 117754. 10.1016/j.envres.2023.117754

Ding, Y.-R., Wang, M.-M., Munipalle, K., Xia, W., Xu, Q., Shen, C., Zhou, T., 2024. Improved exopolysaccharide production by Lactiplantibacillus plantarum Z-1 under hydrogen peroxide stress and its physicochemical properties. Int J Biol Macromol 282, 137215. 10.1016/j.ijbiomac.2024.137215

Dong, C., Beis, K., Nesper, J., Brunkan-LaMontagne, A.L., Clarke, B.R., Whitfield, C., Naismith, J.H., 2006. Wza the translocon for E. coli capsular polysaccharides defines a new class of membrane protein. Nature 444, 226–229. 10.1038/nature05267

Dueholm, M.K.D., Besteman, M., Zeuner, E.J., Riisgaard-Jensen, M., Nielsen, M.E., Vestergaard, S.Z., Heidelbach, S., Bekker, N.S., Nielsen, P.H., 2023. Genetic potential for exopolysaccharide synthesis in activated sludge bacteria uncovered by genome-resolved metagenomics. Water Research 229, 119485. 10.1016/j.watres.2022.119485

Fang, W., Zhang, R., Yang, W., Spanjers, H., Zhang, P., 2024. A novel strategy for waste activated sludge treatment: Recovery of structural extracellular polymeric substances and fermentative production of volatile fatty acids. Water Research 266, 122421. 10.1016/j.watres.2024.122421

Faragò, M., Damgaard, A., Madsen, J.A., Andersen, J.K., Thornberg, D., Andersen, M.H., Rygaard, M., 2021. From wastewater treatment to water resource recovery: Environmental and economic impacts of full-scale implementation. Water Research 204, 117554. 10.1016/j.watres.2021.117554

Felz, S., Vermeulen, P., van Loosdrecht, M.C.M., Lin, Y.M., 2019. Chemical characterization methods for the analysis of structural extracellular polymeric substances (EPS). Water Research 157, 201–208. 10.1016/j.watres.2019.03.068

Fleminger, G., Dayan, A., 2021. The moonlighting activities of dihydrolipoamide dehydrogenase: Biotechnological and biomedical applications. Journal of Molecular Recognition 34, e2924. 10.1002/jmr.2924

González, A., Bellenberg, S., Mamani, S., Ruiz, L., Echeverría, A., Soulère, L., Doutheau, A., Demergasso, C., Sand, W., Queneau, Y., Vera, M., Guiliani, N., 2013. AHL signaling molecules with a large acyl chain enhance biofilm formation on sulfur and metal sulfides by the bioleaching bacterium Acidithiobacillus ferrooxidans. Appl Microbiol Biotechnol 97, 3729–3737. 10.1007/s00253-012-4229-3

Hao, X., Wang, X., Liu, R., Li, S., van Loosdrecht, M.C.M., Jiang, H., 2019. Environmental impacts of resource recovery from wastewater treatment plants. Water Research 160, 268–277. 10.1016/j.watres.2019.05.068

Harrison-McMonagle, P., Denissova, N., Martirnez-Hackert, E., Ebright, R.H., Stock, A.M., 1999. Orientation of OmpR monomers within an OmpR:DNA complex determined by DNA affinity cleaving. Journal of Molecular Biology 285, 555–566. 10.1006/jmbi.1998.2375

Hu, Y.-Q., Wei, W., Gao, M., Zhou, Y., Wang, G.-X., Zhang, Y., 2019. Effect of pure oxygen aeration on extracellular polymeric substances (EPS) of activated sludge treating saline wastewater. Process Safety and Environmental Protection 123, 344–350. 10.1016/j.psep.2019.01.028

Imlay, J.A., 2013. The molecular mechanisms and physiological consequences of oxidative stress: Lessons from a model bacterium. Nat Rev Microbiol 11, 443–454. 10.1038/nrmicro3032

Ionescu, M., Belkin, S., 2009. Overproduction of exopolysaccharides by an Escherichia coli K-12 rpoS mutant in response to osmotic stress. Applied and Environmental Microbiology 75, 483–492. 10.1128/AEM.01616-08

Jang, I.-A., Kim, J., Park, W., 2016. Endogenous hydrogen peroxide increases biofilm formation by inducing exopolysaccharide production in Acinetobacter oleivorans DR1. Sci Rep 6, 21121. 10.1038/srep21121

Ji, J., Qiu, J., Wai, N., Wong, F.-S., Li, Y., 2010. Influence of organic and inorganic flocculants on physical– chemical properties of biomass and membrane-fouling rate. Water Research 44, 1627–1635. 10.1016/j.watres.2009.11.013

Johnson, L.A., Hug, L.A., 2019. Distribution of reactive oxygen species defense mechanisms across domain bacteria. Free Radic Biol Med 140, 93–102. 10.1016/j.freeradbiomed.2019.03.032

Korge, P., Calmettes, G., Weiss, J.N., 2015. Increased reactive oxygen species production during reductive stress: The roles of mitochondrial glutathione and thioredoxin reductases. Biochimica et Biophysica Acta (BBA) - Bioenergetics 1847, 514–525. 10.1016/j.bbabio.2015.02.012

Korshunov, S., Imlay, J.A., 2024. Antioxidants are ineffective at quenching reactive oxygen species inside bacteria and should not be used to diagnose oxidative stress. Molecular Microbiology 122, 113–128. 10.1111/mmi.15286

Kroiss, H., 2004. What is the potential for utilizing the resources in sludge? Water Science and Technology 49, 1–10. 10.2166/wst.2004.0595

Li, J., Hao, X., Gan, W., van Loosdrecht, M.C.M., Wu, Y., 2021. Recovery of extracellular biopolymers from conventional activated sludge: Potential, characteristics and limitation. Water Research 205, 117706. 10.1016/j.watres.2021.117706

Li, W.-W., Yu, H.-Q., Rittmann, B.E., 2015. Chemistry: Reuse water pollutants. Nature 528, 29–31. 10.1038/528029a

Li, X.Y., Yang, S.F., 2007. Influence of loosely bound extracellular polymeric substances (EPS) on the flocculation, sedimentation and dewaterability of activated sludge. Water Research 41, 1022–1030. 10.1016/j.watres.2006.06.037

Liu, X., Huang, D., Zhu, C., Zhu, F., Zhu, X., Zhou, D., 2024. Production of reactive oxygen species during redox manipulation and its potential impacts on activated sludge wastewater treatment processes. Environ Sci Technol 58, 23042–23052. 10.1021/acs.est.4c11301

Mao, Z., Shin, H.-D., Chen, R.R., 2006. Engineering the E. coli UDP-glucose synthesis pathway for oligosaccharide synthesis. Biotechnology Progress 22, 369–374. 10.1021/bp0503181

Messner, K.R., Imlay, J.A., 1999. The identification of primary sites of superoxide and hydrogen peroxide formation in the aerobic respiratory chain and sulfite reductase complex of Escherichia coli. Journal of Biological Chemistry 274, 10119–10128. 10.1074/jbc.274.15.10119

Prigent-Combaret, C., Brombacher, E., Vidal, O., Ambert, A., Lejeune, P., Landini, P., Dorel, C., 2001. Complex regulatory network controls initial adhesion and biofilm formation in Escherichia coli via regulation of the csgD gene. J Bacteriol 183, 7213–7223. 10.1128/JB.183.24.7213-7223.2001

Ravindra Kumar, S., Imlay, J.A., 2013. How Escherichia coli tolerates profuse hydrogen peroxide formation by a catabolic pathway. Journal of Bacteriology 195, 4569–4579. 10.1128/jb.00737-13

Salama, Y., Chennaoui, M., Sylla, A., Mountadar, M., Rihani, M., Assobhei, O., 2016. Characterization, structure, and function of extracellular polymeric substances (EPS) of microbial biofilm in biological wastewater treatment systems: A review. Desalination and Water Treatment 57, 16220–16237. 10.1080/19443994.2015.1077739

Schmid, J., 2018. Recent insights in microbial exopolysaccharide biosynthesis and engineering strategies. Current Opinion in Biotechnology, Chemical Biotechnology • Pharmaceutical Biotechnology 53, 130–136. 10.1016/j.copbio.2018.01.005

Seaver, L.C., Imlay, J.A., 2004. Are respiratory enzymes the primary sources of intracellular hydrogen peroxide? Journal of Biological Chemistry 279, 48742–48750. 10.1074/jbc.M408754200

Seo, S.W., Gao, Y., Kim, D., Szubin, R., Yang, J., Cho, B.-K., Palsson, B.O., 2017. Revealing genome-scale transcriptional regulatory landscape of OmpR highlights its expanded regulatory roles under osmotic stress in Escherichia coli K-12 MG1655. Sci Rep 7, 2181. 10.1038/s41598-017-02110-7

Sheng, G.-P., Yu, H.-Q., Li, X.-Y., 2010. Extracellular polymeric substances (EPS) of microbial aggregates in biological wastewater treatment systems: A review. Biotechnology Advances 28, 882–894. 10.1016/j.biotechadv.2010.08.001

Shi, Y., Huang, J., Zeng, G., Gu, Y., Chen, Y., Hu, Y., Tang, B., Zhou, J., Yang, Y., Shi, L., 2017. Exploiting extracellular polymeric substances (EPS) controlling strategies for performance enhancement of biological wastewater treatments: An overview. Chemosphere 180, 396–411. 10.1016/j.chemosphere.2017.04.042

Shi, Y., Xing, S., Wang, X., Wang, S., 2013. Changes of the reactor performance and the properties of granular sludge under tetracycline (TC) stress. Bioresource Technology 139, 170–175. 10.1016/j.biortech.2013.03.037

Shu, C.-H., Lung, M.-Y., 2004. Effect of pH on the production and molecular weight distribution of exopolysaccharide by Antrodia camphorata in batch cultures. Process Biochemistry 39, 931–937. 10.1016/S0032-9592(03)00220-6

Sies, H., 2021. Oxidative eustress: On constant alert for redox homeostasis. Redox Biol 41, 101867. 10.1016/j.redox.2021.101867

Tan, C.H., Koh, K.S., Xie, C., Tay, M., Zhou, Y., Williams, R., Ng, W.J., Rice, S.A., Kjelleberg, S., 2014. The role of quorum sensing signalling in EPS production and the assembly of a sludge community into aerobic granules. The ISME Journal 8, 1186–1197. 10.1038/ismej.2013.240

van Loosdrecht, M.C.M., Brdjanovic, D., 2014. Anticipating the next century of wastewater treatment. Science 344, 1452–1453. 10.1126/science.1255183

Wingender, J., Neu, T.R., Flemming, H.-C., 1999. What are Bacterial Extracellular Polymeric Substances?, in: Wingender, J., Neu, T.R., Flemming, H.-C. (Eds.), Microbial Extracellular Polymeric Substances: Characterization, Structure and Function. Springer, Berlin, Heidelberg, pp. 1–19. 10.1007/978-3-642-60147-7_1

Yang, H., Zhou, Y., Luo, Q., Zhu, C., Fang, B., 2023. L-leucine increases the sensitivity of drug-resistant Salmonella to sarafloxacin by stimulating central carbon metabolism and increasing intracellular reactive oxygen species level. Front. Microbiol. 14. 10.3389/fmicb.2023.1186841

Ye, F., Ye, Y., Li, Y., 2011. Effect of C/N ratio on extracellular polymeric substances (EPS) and physicochemical properties of activated sludge flocs. Journal of Hazardous Materials 188, 37–43. 10.1016/j.jhazmat.2011.01.043

Yuniarto, A., Noor, Z.Z., Ujang, Z., Olsson, G., Aris, A., Hadibarata, T., 2013. Bio-fouling reducers for improving the performance of an aerobic submerged membrane bioreactor treating palm oil mill effluent. Desalination 316, 146–153. 10.1016/j.desal.2013.02.002

Zhang, L., Alfano, J.R., Becker, D.F., 2015. Proline metabolism increases katG expression and oxidative stress resistance in Escherichia coli. Journal of Bacteriology 197, 431–440. 10.1128/jb.02282-14

Zhou, X., Manna, B., Lyu, B., Lear, G., Kingsbury, J.M., Singhal, N., 2025. Resource recovery from wastewater by directing microbial metabolism toward production of value-added biochemicals. Bioresource Technology 419, 132061. 10.1016/j.biortech.2025.132061

