## Supplementary for "Linking Oxygen-Induced Oxidative Stress to Resource Recovery by Enhancing the Production of Extracellular Polymeric Substances in Activated Sludge Microbial Communities"

34 pages, 27 figures and 4 tables

**Table S3.** Summary of the 60 metabolism-associated flavoenzymes and their corresponding electron-donating substrates under different aeration conditions. 12

**Figure S22.** Sankey visualization of microbial taxa contributing to K00318 (proline dehydrogenase) under CP conditions. 26
**Figure S23.** Sankey visualization of microbial taxa contributing to K00318 (proline dehydrogenase) under CA conditions. 27
**Figure S24.** Sankey visualization of microbial taxa contributing to K00318 (proline dehydrogenase) under IP conditions. 28
**Figure S25.** Sankey visualization of microbial taxa contributing to K00263 (leucine dehydrogenase) under CP conditions. 29
**Figure S26.** Sankey visualization of microbial taxa contributing to K00263 (leucine dehydrogenase) under CA conditions. 30
**Figure S27.** Sankey visualization of microbial taxa contributing to K00263 (leucine dehydrogenase) under IP conditions. 31
**Table S4.** Taxonomic origins of six ROS-generating enzymes identified in the metaproteomic dataset. 31
**S2.3. Potential yield and economic value of EPS enhancement under full-scale conditions** 32
**References** 34

### S1. Supplementary methods

#### S1.1. Bioreactor operation and performance

**Table S1.** Composition of the trace element solution

| Chemical | Concentration (g/L) |
|---|---|
| EDTA | 2.50 |
| $ZnSO_4 \cdot 7H_2O$ | 1.10 |
| $CoCl_2 \cdot 6H_2O$ | 0.80 |
| $MnCl_2 \cdot 4H_2O$ | 2.55 |
| $MgSO_4 \cdot 7H_2O$ | 20.0 |
| $CuSO_4 \cdot 5H_2O$ | 0.86 |
| $(NH_4)_6Mo_7O_{24} \cdot 4H_2O$ | 0.07 |
| $CaCl_2 \cdot 2H_2O$ | 2.75 |
| $FeSO_4 \cdot 7H_2O$ | 2.57 |

**Table S2.** Composition of the concentrated artificial wastewater

| Chemical | Concentration |
|---|---|
| Methanol (mL/L) | 26.94 |
| $NH_4Cl$ (g/L) | 24.46 (IP); 30.57 (CP or CA) |
| $KH_2PO_4$ (g/L) | 2.76 |
| $K_2HPO_4$ (g/L) | 2.76 |
| $NaHCO_3$ (g/L) | 76.80 |

#### S1.2. The three aeration conditions

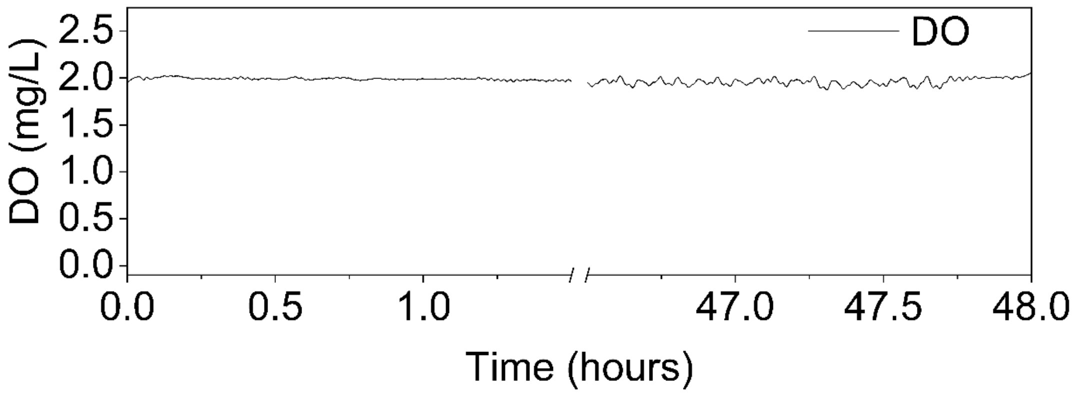

**Figure S1.** Dissolved oxygen concentrations were controlled under Constant Aeration (CA) to maintain a stable DO level at 2.0 ± 0.2 mg/L.

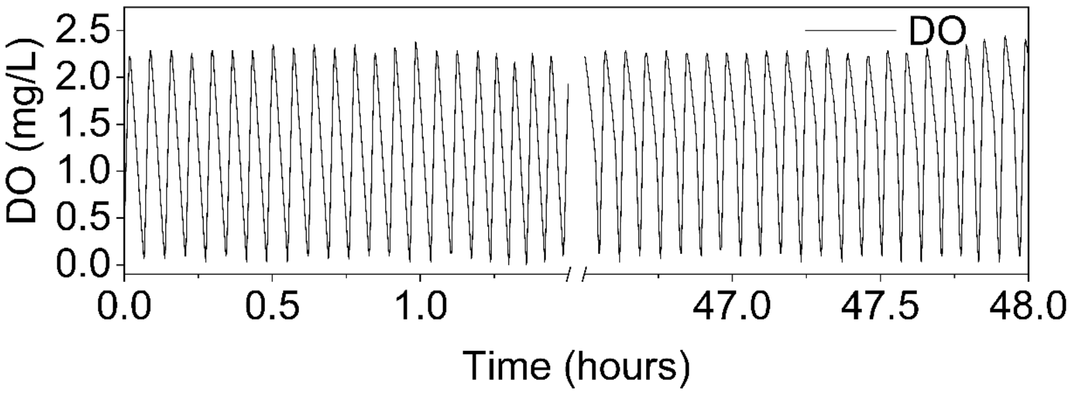

**Figure S2.** Dissolved oxygen concentrations under Continuous Perturbation (CP) were controlled to continuously fluctuate between 0.1 and 2.0 mg/L without intermission.

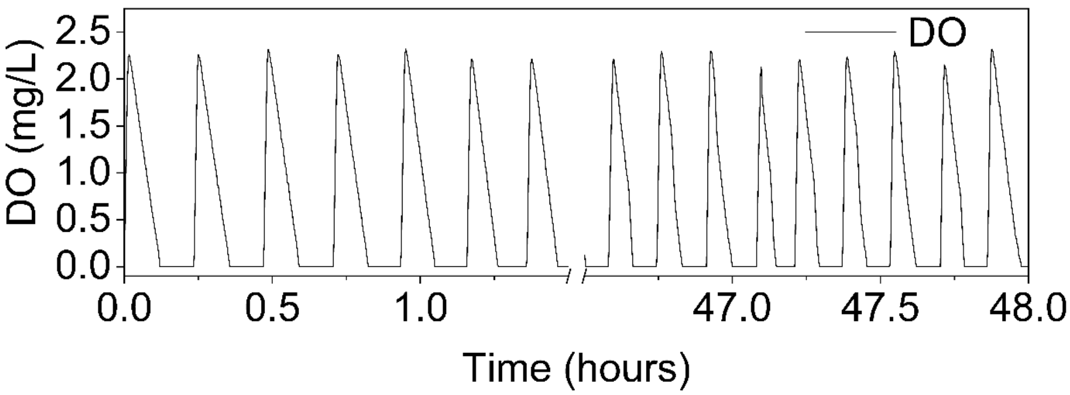

**Figure S3.** Dissolved oxygen concentrations under Intermittent Perturbation (IP) were controlled to fluctuate between 0 and 2.0 mg/L, with equal durations of aerobic and anoxic phases.

**S1.3. DNA extraction, metagenomic sequencing, and bioinformatic analysis**

Genomic DNA was extracted from 10 mL of sludge samples using the DNeasy PowerSoil Kit (Qiagen, Germany), following the manufacturer's recommended protocol. The purified DNA was stored at –20 °C until further analysis. Metagenomic sequencing was performed by the Auckland Genomics Centre. Prior to sequencing, DNA concentration and quality were assessed, followed by PCR amplification and indexing for library preparation. After library quality control and normalization, a final pooling step was conducted using a bioanalyzer. Sequencing was carried out on the Illumina HiSeq platform, producing approximately 400 million paired-end reads (2 × 150 bp). Raw sequencing data underwent quality filtering using Trimmomatic v0.39 (Bolger et al., 2014), applying the following parameters: LEADING:3, TRAILING:3, SLIDINGWINDOW:10:15, and MINLEN:50, with adapter trimming based on the TruSeq3-PE-2.fa file. Metagenomic assembly, taxonomic assignment, and functional annotation were performed using the SqueezeMeta v1.5.2 pipeline (Tamames and Puente-Sánchez, 2019). Co-assembly was executed using MEGAHIT v1.2.9 (Li et al., 2015), and contigs shorter than 200 bp were excluded using PRINSEQ v0.20.4 (Schmieder and Edwards, 2011). Ribosomal RNA prediction and open reading frame (ORF) identification were conducted with Barrnap v0.9 (Seemann, 2014) and Prodigal v2.6.3 (Hyatt et al., 2010), respectively. Taxonomic classification was based on the NCBI GenBank nr database, using identity thresholds of 85%, 60%, 55%, 50%, 46%, 42%, and 40% for species to superkingdom levels., respectively. Functional annotation was performed against the KEGG database using DIAMOND v2.0.14 (Buchfink et al., 2015), with a maximum e-value of $1 \times 10^{-3}$ and a minimum sequence identity of 50%. Read alignment to assembled contigs was conducted using Bowtie2 v2.3.4.1 (Langmead and Salzberg, 2012). Abundances of functional genes were normalized and expressed in transcripts per million (TPM). All metagenomic outputs were further processed and visualized using the SQMtools R package v1.6.3 (Puente-Sánchez et al., 2020).

To identify the taxonomic contributors to specific KEGG functions, all predicted ORFs were first taxonomically classified based on sequence homology against the GenBank nr database using a Last Common Ancestor (LCA) approach, applying identity thresholds of 85%, 60%, 55%, 50%, 46%, 42%, and 40% for species to superkingdom levels, respectively. Functional annotation was performed against the KEGG database as described above. Genes sharing the same functional annotation were subsetted using the subsetFun() function of the SQMtools package. The taxonomic affiliation of these functional genes was assigned based on the taxonomic classification of their parent contigs.

All metagenomic sequencing data generated in this study have been deposited in the European Nucleotide Archive (ENA) under the project accession number PRJEB74089.

**S1.4. Protein extraction, metaproteomic analysis, and data processing**

At the end of the reaction, 5 mL of mixed liquor was sampled from the bioreactor. Biomass was collected by centrifugation at 6,113 × g for 5 min at 4 °C, followed by washing and pelleting. The resulting pellets were rapidly frozen in liquid nitrogen and stored at −80 °C until protein extraction. For lysis, frozen pellets were resuspended in lysis buffer and disrupted on ice via sonication for 20 cycles. Cell lysates were clarified by centrifugation at 17,467 × g for 30 min at 4 °C.

Proteins were precipitated by adding 20% trichloroacetic acid in acetone, vortexed briefly, and incubated on ice. Following centrifugation, the supernatant was discarded, and the protein pellet was washed with pure acetone and then with 80% aqueous acetone, with centrifugation steps in between. After air-drying, the protein pellet was dissolved in 50 mM Tris buffer. Protein purification was conducted using SpeedBead Carboxylate-Modified Magnetic Particles E3 and E7 (Sera-Mag, USA), and total protein content was quantified using the EZQ Protein Assay Kit (Invitrogen, USA). Samples were then subjected to standard reduction, alkylation, and enzymatic digestion with trypsin.

Post-digestion, peptides were diluted, filtered, and purified using solid-phase extraction. The eluates were concentrated by vacuum centrifugation, and 10 μL aliquots were injected for LC-MS/MS analysis. Chromatographic separation and desalting were performed on a NanoLC 400 UPLC system (Eksigent, USA), while mass spectrometry was carried out using a TripleTOF 6600 Q-TOF instrument (Sciex, USA).

Protein identification was performed using MetaProteomeAnalyzer v3.4 (Muth et al., 2015), with peptide-spectrum matching conducted via the X! Tandem (Xu et al., 2013) search engine. Searches were conducted against a custom protein database derived from the sample-specific metagenomic data. Search parameters included a precursor and fragment mass tolerance of 15 ppm, fixed cysteine alkylation with iodoacetamide, and trypsin digestion allowing up to one missed cleavage. A 1% false discovery rate (FDR) threshold was applied to define global protein groups. Final protein grouping and clustering were performed within MetaProteomeAnalyzer.

In this study, flavoenzymes were identified by referencing the gene ID information of prokaryotic flavoenzymes from the UniProt database (The UniProt Consortium, 2025). The corresponding metabolic pathways of the detected proteins were annotated according to KEGG pathway classifications (Kanehisa et al., 2025).

The metaproteomics data has been submitted to ProteomeXchange Consortium (Perez-Riverol et al., 2022) with the dataset identifier PXD044490.

**S1.5. Intracellular Metabolite Extraction and GC-MS analysis, and data processing**

Intracellular metabolites were profiled using a gas chromatography–mass spectrometry (GC-MS) platform, with three biological replicates and two technical replicates analyzed for each experimental condition. For each sample, 10 mL of activated sludge was centrifuged, and the supernatant was discarded. The resulting pellet was rapidly quenched in liquid nitrogen and stored at –80 °C until extraction. Metabolites were extracted by adding 2.5 mL of chilled methanol–water solution along with 0.2 µmol of 2,3,3,3-$d_4$-alanine as an internal standard. Intracellular compounds were released via three freeze–thaw cycles followed by vigorous shaking. The mixture was centrifuged at 22,707 × g for 15 min at –20 °C to collect the supernatant. An additional 2.5 mL of cold methanol–water solution was added to the pellet for further extraction, and the supernatants were pooled.

The metabolite extract was reconstituted in 400 µL of 1 M sodium hydroxide, followed by the addition of 68 µL pyridine and 334 µL methanol. Methyl chloroformate (40 µL) was added twice, each followed by 30 s of vortexing. Subsequently, 400 µL chloroform was introduced and vortexed for another 10 s. The mixture was then supplemented with 800 µL of 50 mM sodium bicarbonate, vortexed briefly, and centrifuged to separate the phases. The aqueous layer was discarded, and any residual moisture in the chloroform phase was removed using anhydrous sodium sulfate.

Metabolite derivatization products were analyzed using an Agilent GC7890 gas chromatograph coupled to an MSD597 mass spectrometer, equipped with a ZB-1701 capillary column (30 m × 250 µm i.d. × 0.15 µm film thickness) and a 5 m guard column (Phenomenex, Torrance, CA, USA). Chromatographic deconvolution was performed using AMDIS software (NIST, Boulder, CO, USA), and metabolite identification was achieved by comparing spectra against an in-house mass spectral library (Smart et al., 2010). Raw data were filtered using the MassOmics R package, with peak intensities normalized to the internal standard (Guo et al., 2021). Blank-corrected relative abundances of intracellular metabolites were calculated over the 48-hour period.

The mass spectrometry metabolomics data have been deposited to MetaboLights (Haug et al., 2020) with the dataset identifier MTBLS8331.

**S2. Results**

**S2.1. Reactive oxygen and nitrogen species and antioxidant system analysis**

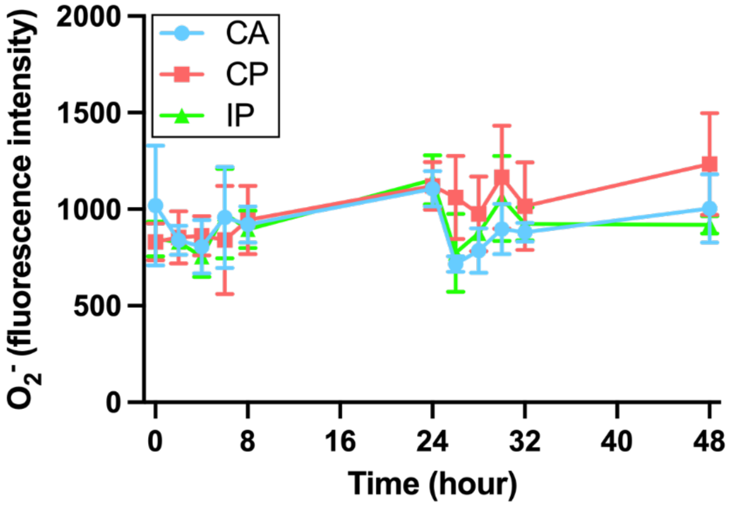

**Figure S4.** The fluorescence intensities of $O_2^{\bullet-}$ under different aeration conditions: constant aeration (CA), continuous perturbation (CP), and intermittent perturbation (IP). *Error bars represent SD (n = 3 biological replicates); significance tested by repeated-measures ANOVA followed by Tukey's multiple comparisons test*.

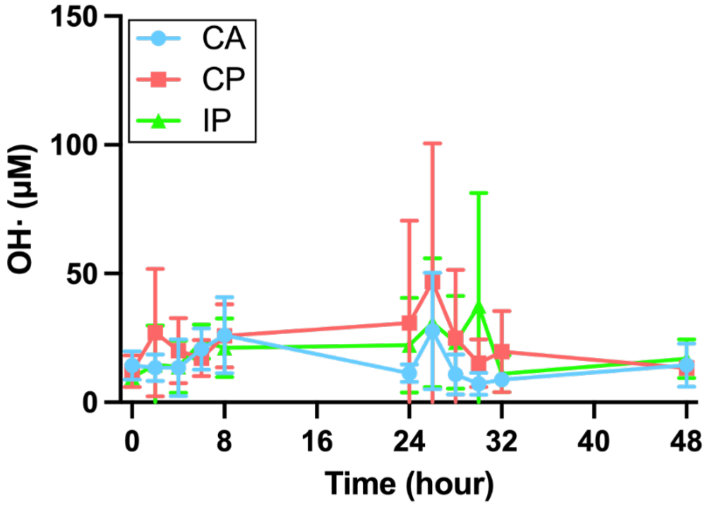

**Figure S5.** The concentration of OH· under different aeration conditions: constant aeration (CA), continuous perturbation (CP), and intermittent perturbation (IP). *Error bars represent SD (n = 3 biological replicates); significance tested by repeated-measures ANOVA followed by Tukey's multiple comparisons test.*

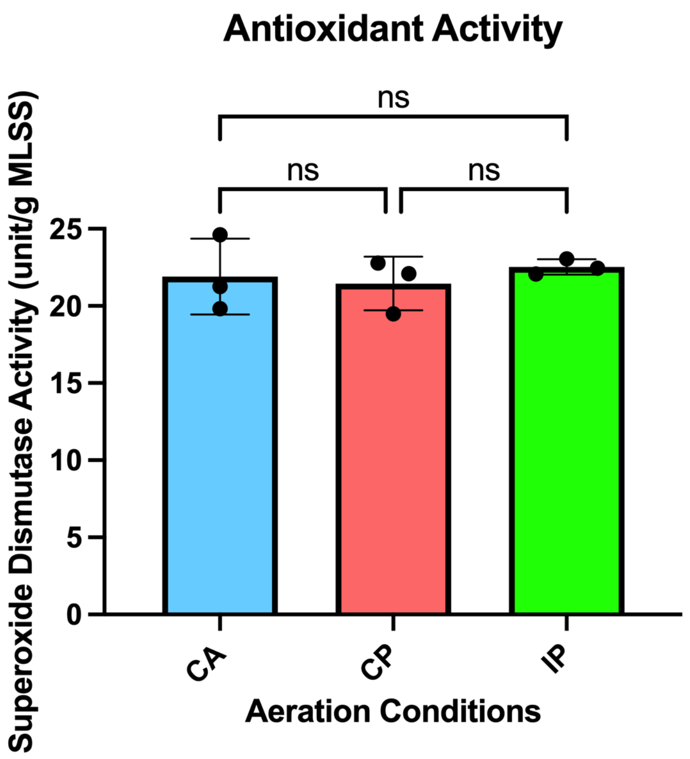

**Figure S6.** The activity of superoxide dismutase following incubation for 48 hours under different aeration conditions: constant aeration (CA), continuous perturbation (CP), and intermittent perturbation (IP). *Error bars represent SD (n = 3 biological replicates); significance tested by one-way ANOVA.*

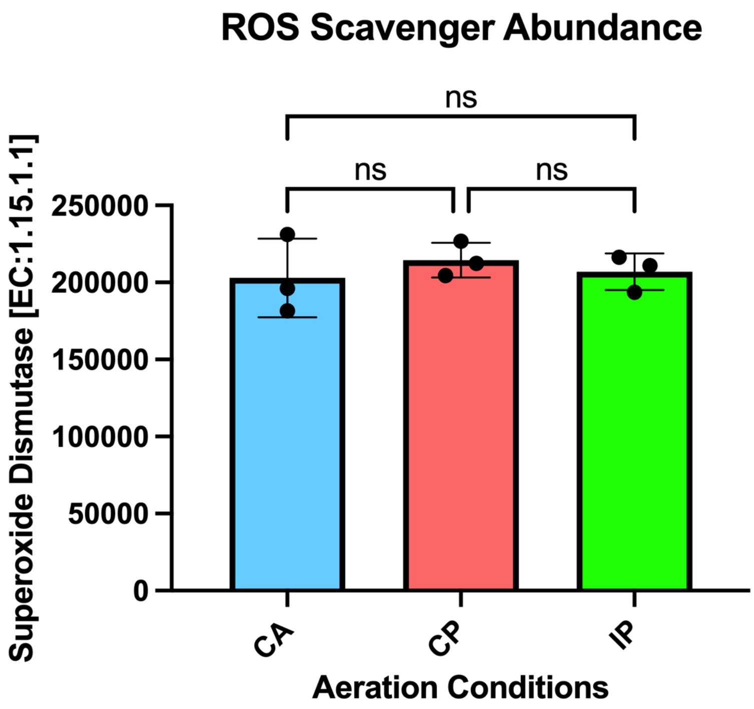

**Figure S7.** The abundance of superoxide dismutase following incubation for 48 hours under different aeration conditions: constant aeration (CA), continuous perturbation (CP), and intermittent perturbation (IP). *Error bars represent SD (n = 3 biological replicates); significance tested by one-way ANOVA.*

### S2.2. Microbial contributors to endogenous ROS production

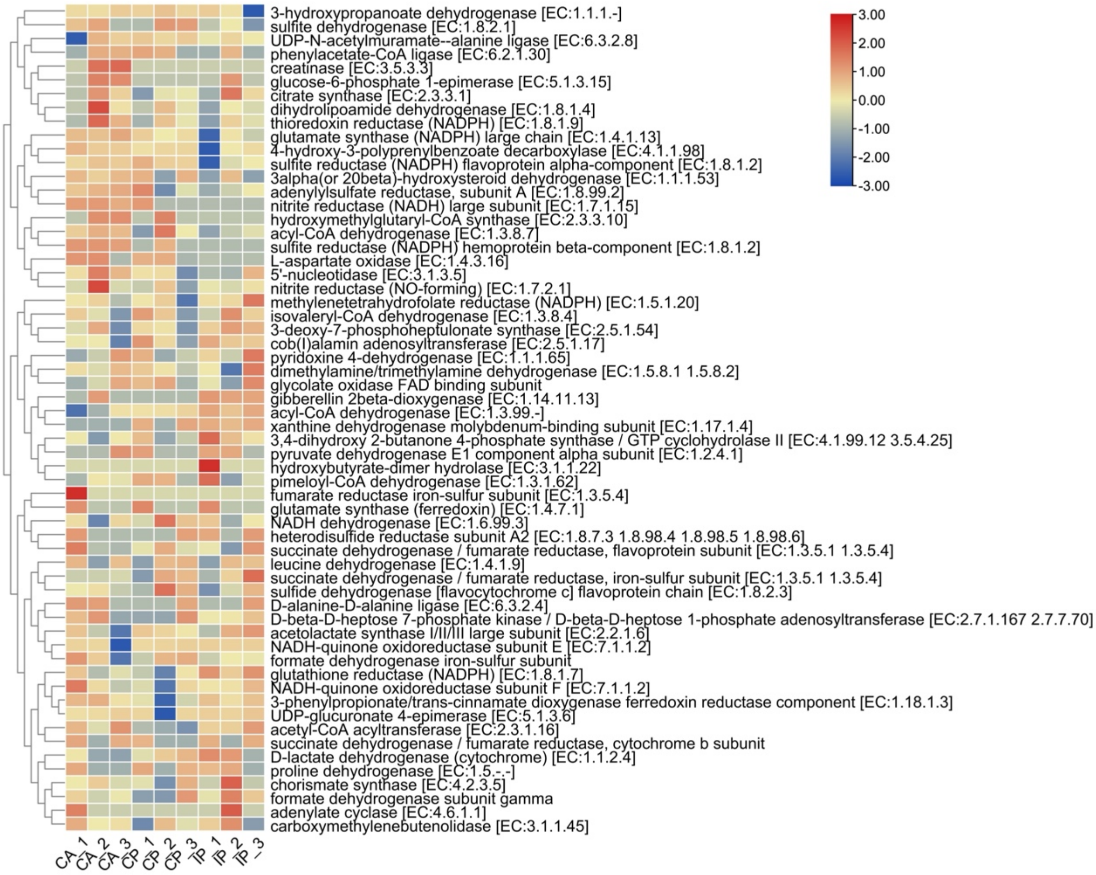

**Figure S8.** Heatmap showing metabolism-related flavoenzymes identified in the metaproteomic dataset. Only pathways classified under the KEGG Level 1 category "metabolism" are included.

**Table S3.** Summary of the 60 metabolism-associated flavoenzymes and their corresponding electron-donating substrates under different aeration conditions.
[This table is provided in Excel format to facilitate pathway-level interpretation of ROS-producing enzymes.]

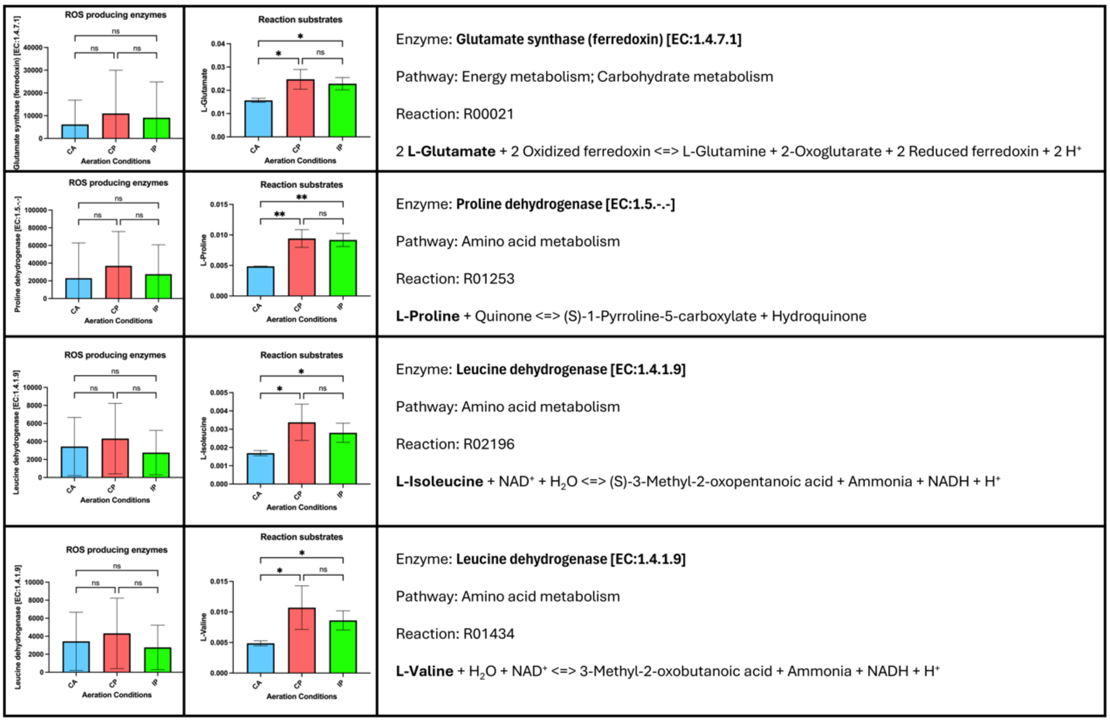

**Figure S9.** Relative protein abundance of three ROS-related flavoenzymes, including glutamate synthase (ferredoxin), proline dehydrogenase, and leucine dehydrogenase, and corresponding electron-donating substrate concentrations under CA, CP, and IP. *Error bars represent standard deviation; n = 3 biological replicates.*

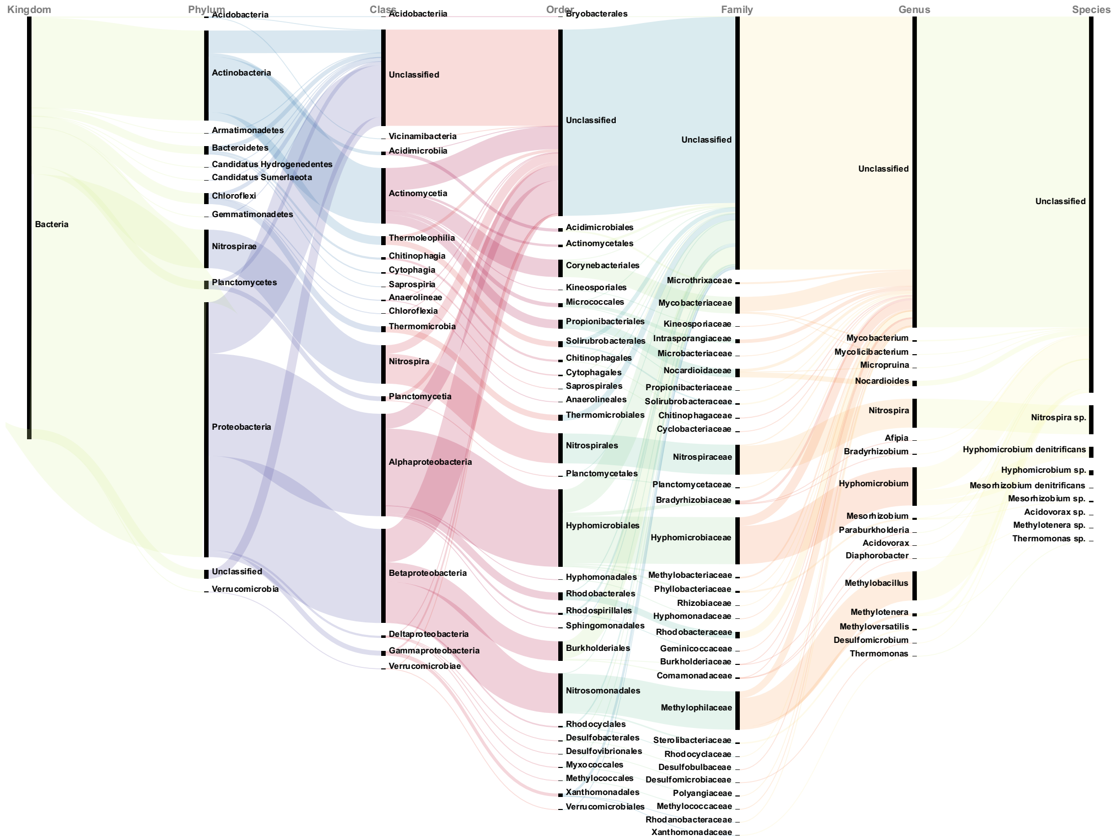

**Figure S10.** Sankey visualization of microbial taxa contributing to K00265 (glutamate synthase, NADPH) under CP conditions.

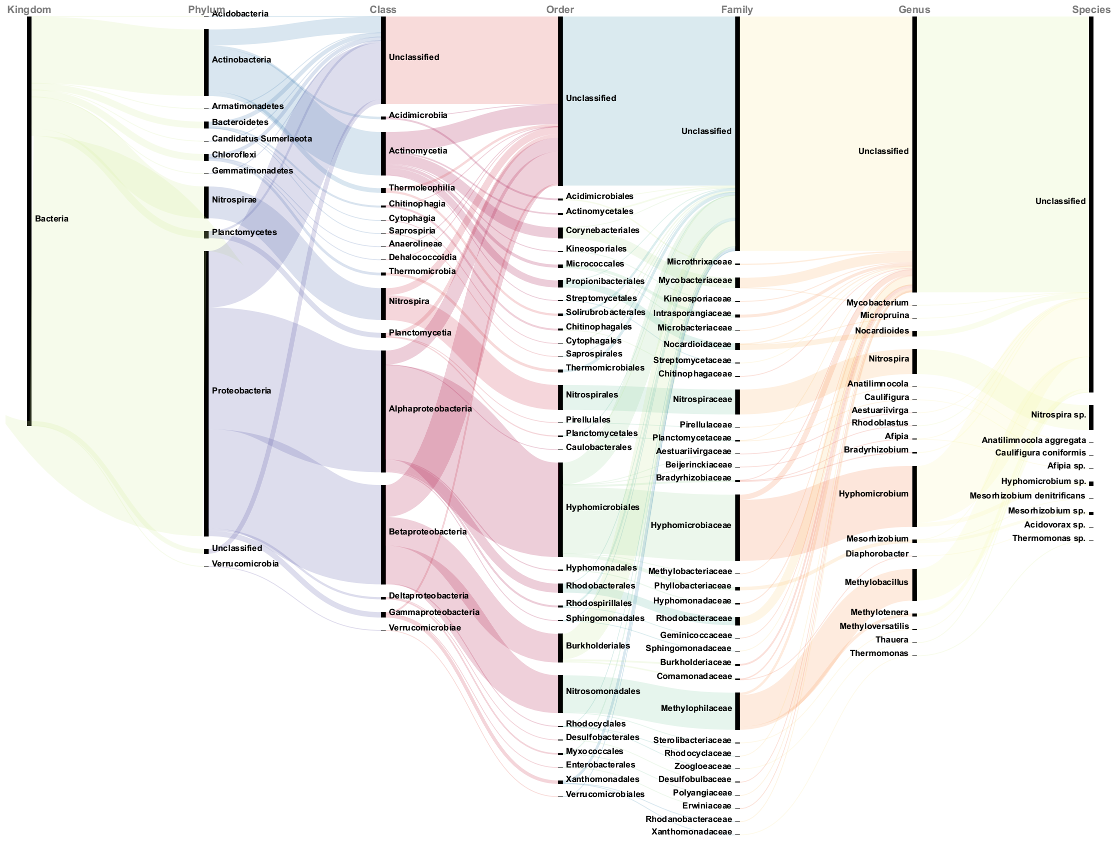

**Figure S11.** Sankey visualization of microbial taxa contributing to K00265 (glutamate synthase, NADPH) under CA conditions.

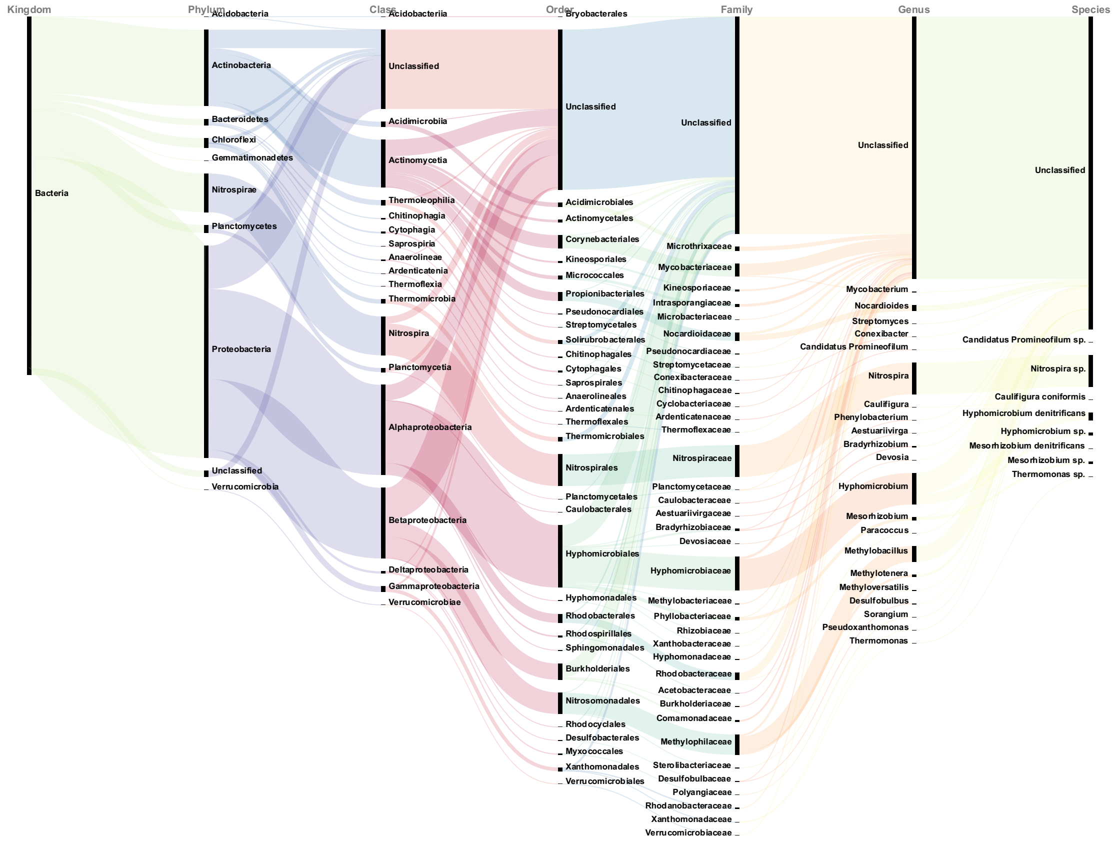

**Figure S12.** Sankey visualization of microbial taxa contributing to K00265 (glutamate synthase, NADPH) under IP conditions.

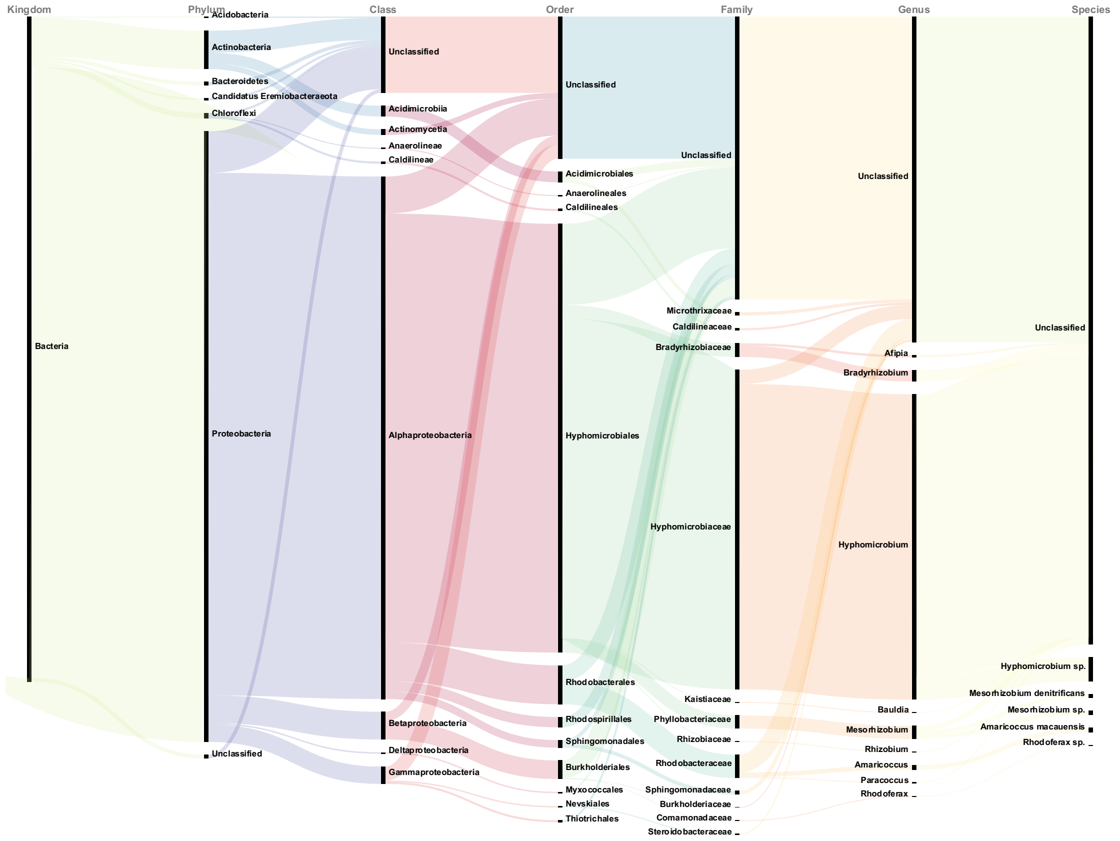

**Figure S13.** Sankey visualization of microbial taxa contributing to K00383 (glutathione reductase) under CP conditions.

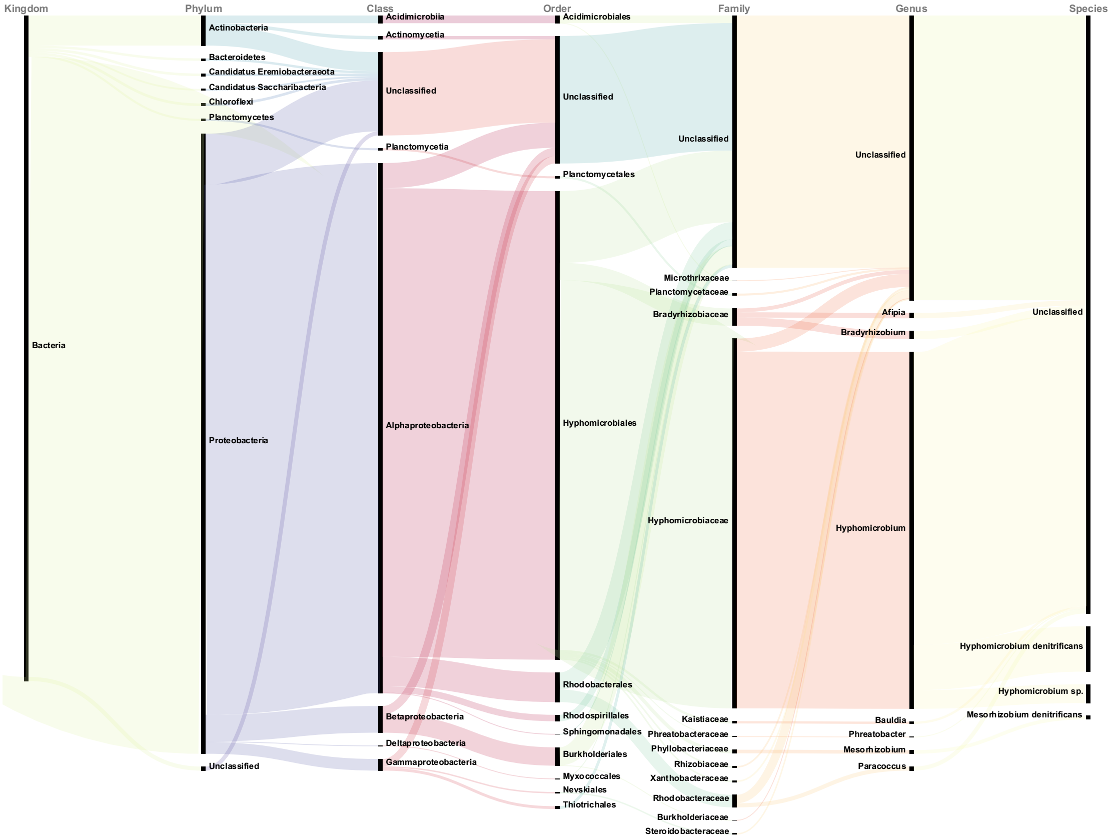

**Figure S14.** Sankey visualization of microbial taxa contributing to K00383 (glutathione reductase) under CA conditions.

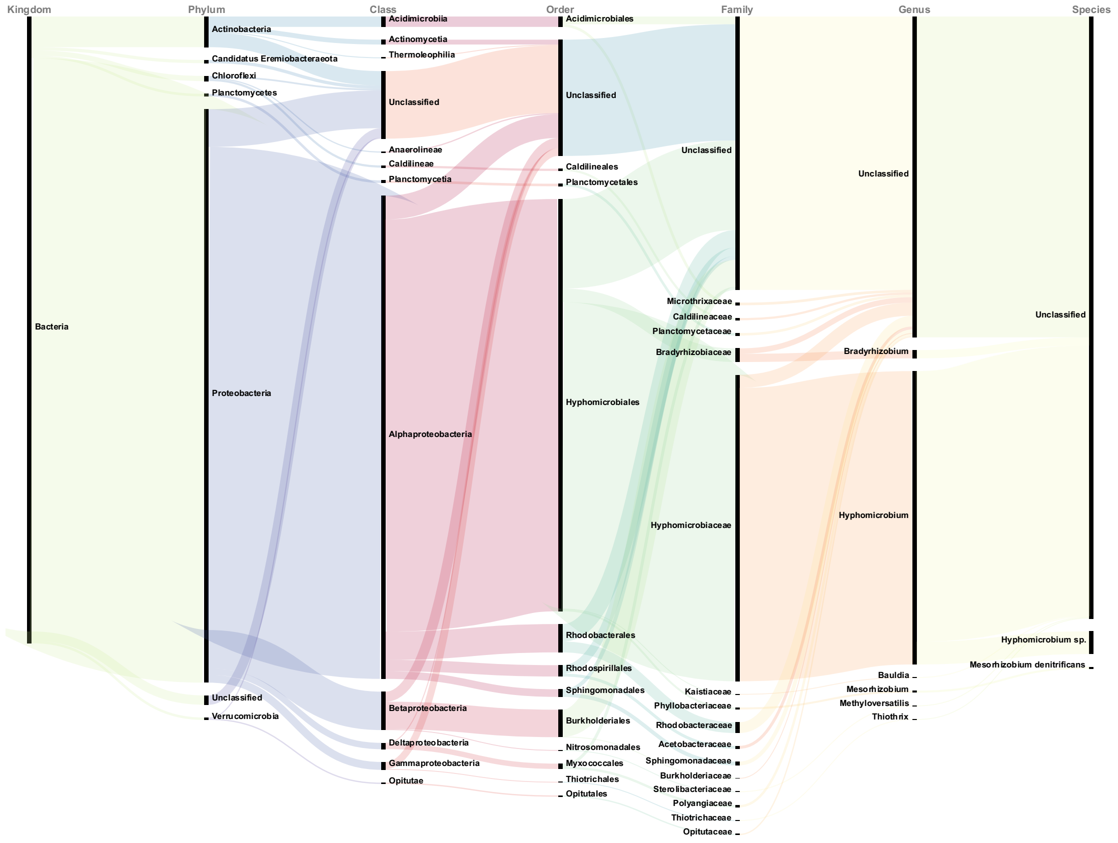

**Figure S15.** Sankey visualization of microbial taxa contributing to K00383 (glutathione reductase) under IP conditions.

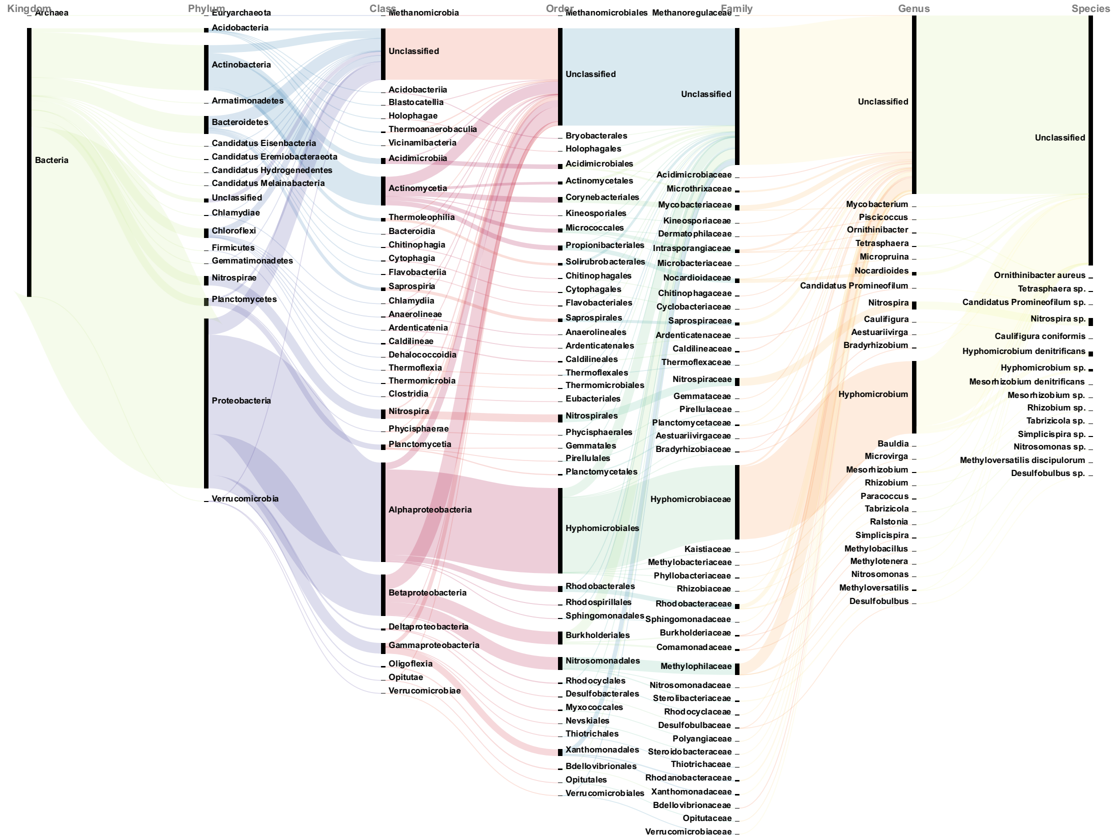

**Figure S16.** Sankey visualization of microbial taxa contributing to K00382 (dihydrolipoamide dehydrogenase) under CP conditions.

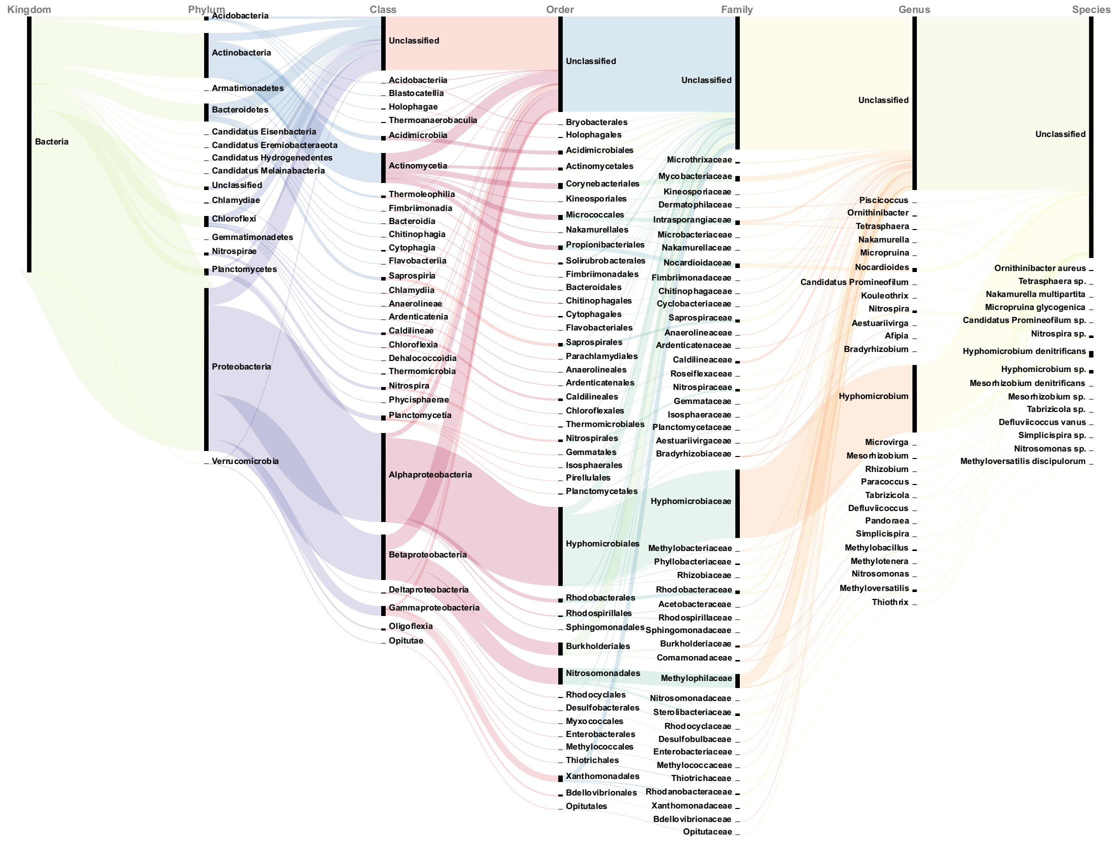

**Figure S17.** Sankey visualization of microbial taxa contributing to K00382 (dihydrolipoamide dehydrogenase) under CA conditions.

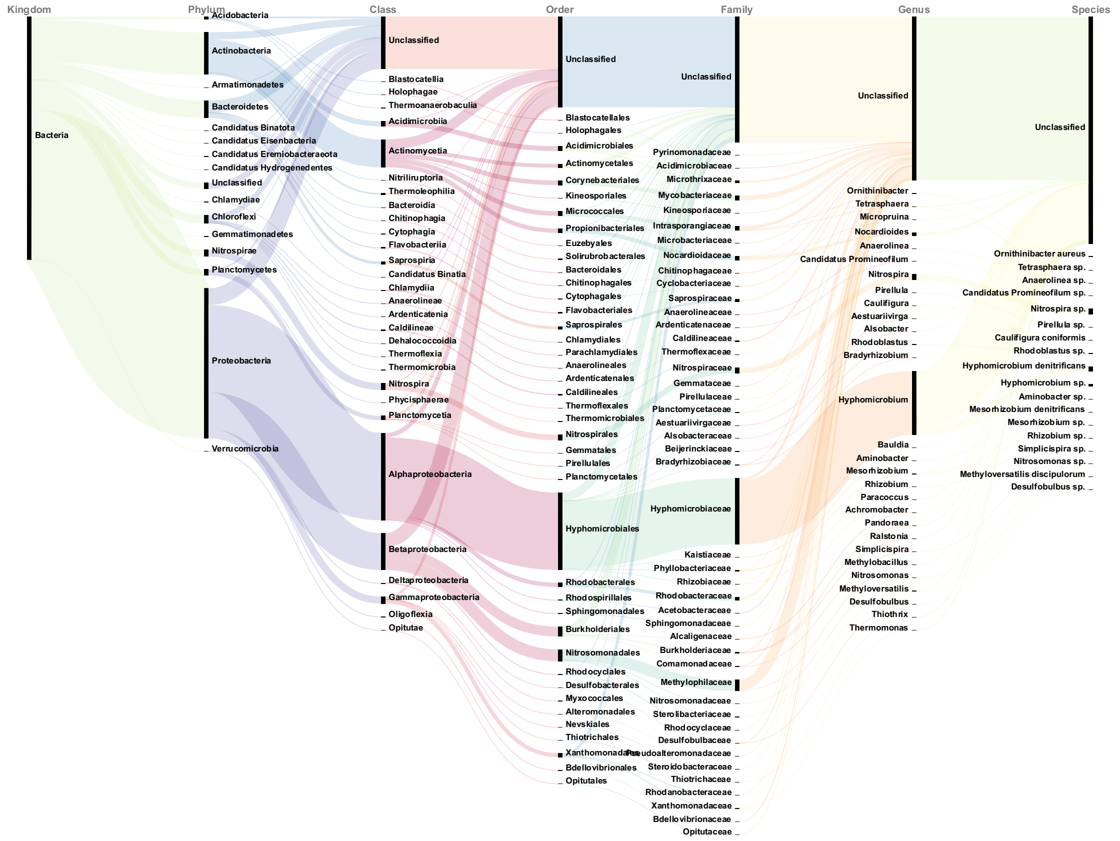

**Figure S18.** Sankey visualization of microbial taxa contributing to K00382 (dihydrolipoamide dehydrogenase) under IP conditions.

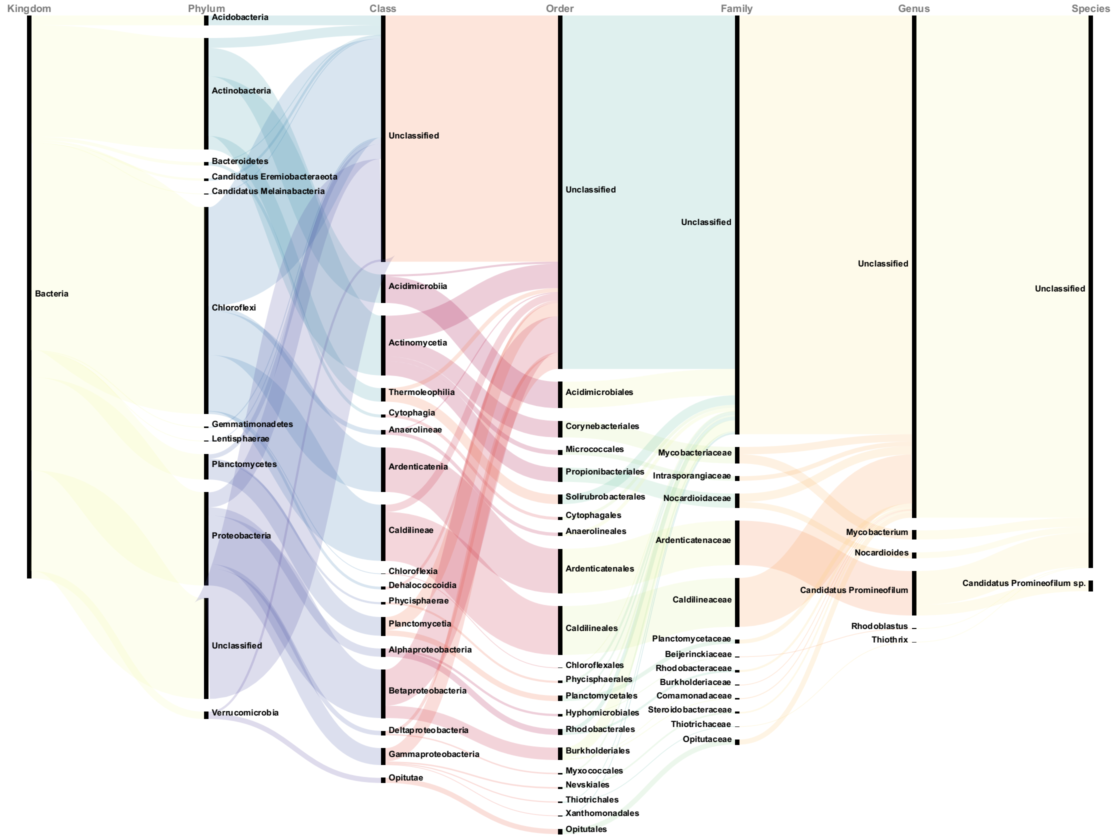

**Figure S19.** Sankey visualization of microbial taxa contributing to K00284 (glutamate synthase, ferredoxin) under CP conditions.

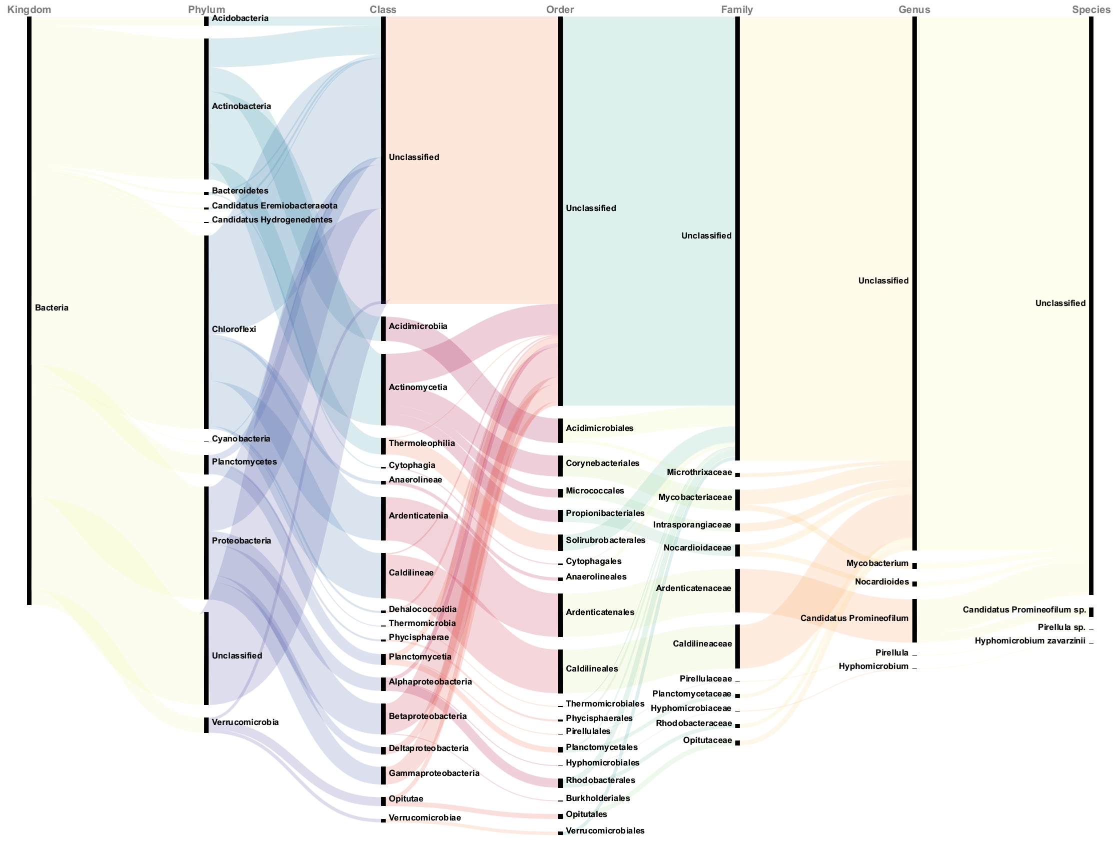

**Figure S20.** Sankey visualization of microbial taxa contributing to K00284 (glutamate synthase, ferredoxin) under CA conditions.

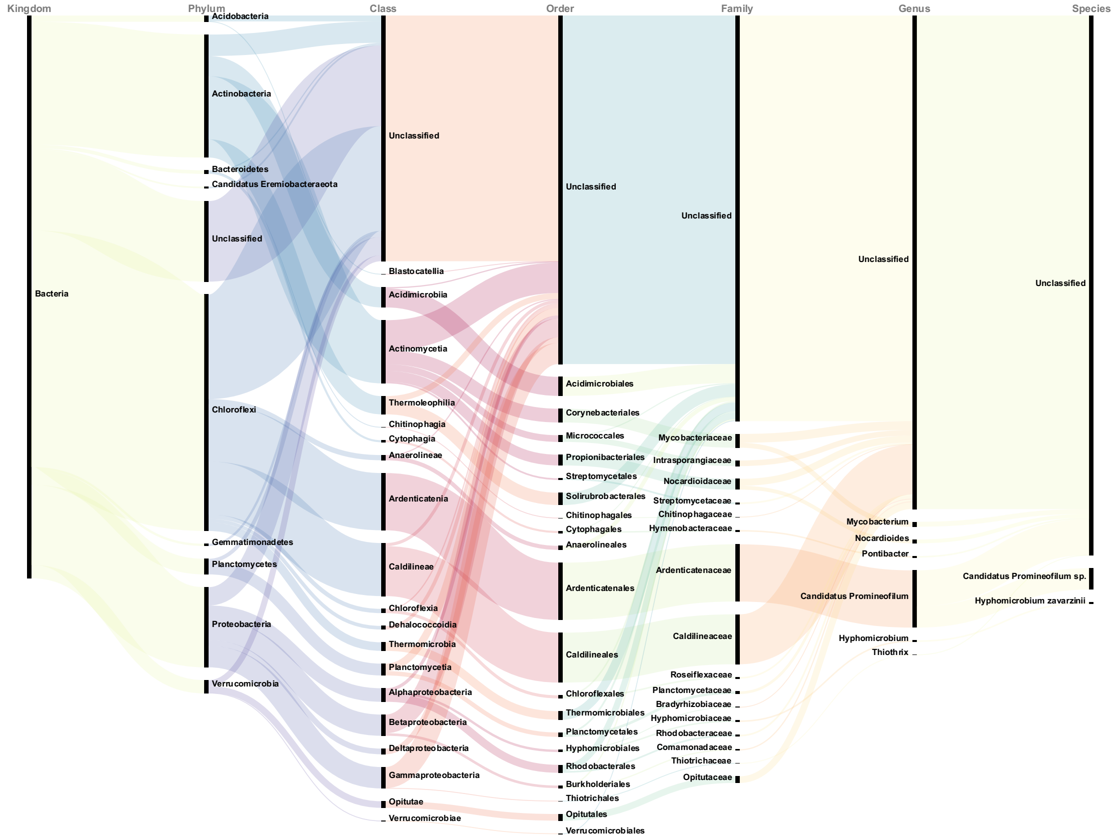

**Figure S21.** Sankey visualization of microbial taxa contributing to K00284 (glutamate synthase, ferredoxin) under IP conditions.

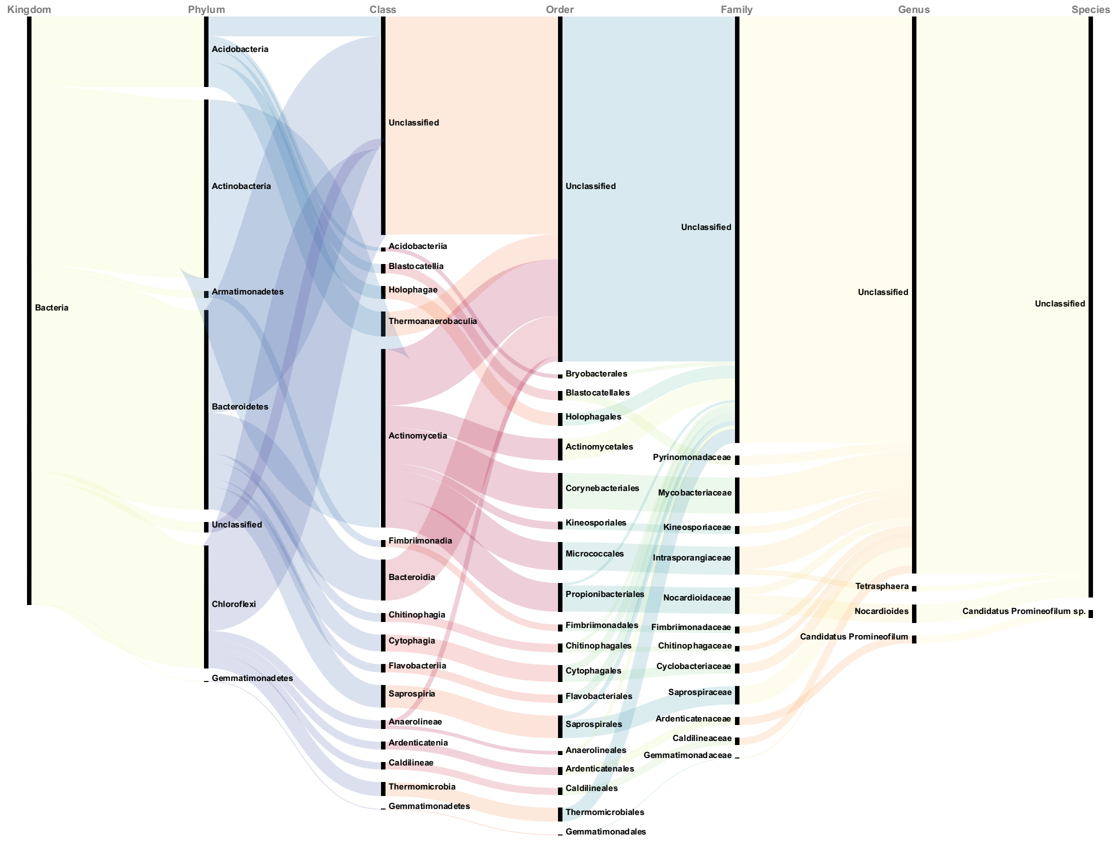

**Figure S22.** Sankey visualization of microbial taxa contributing to K00318 (proline dehydrogenase) under CP conditions.

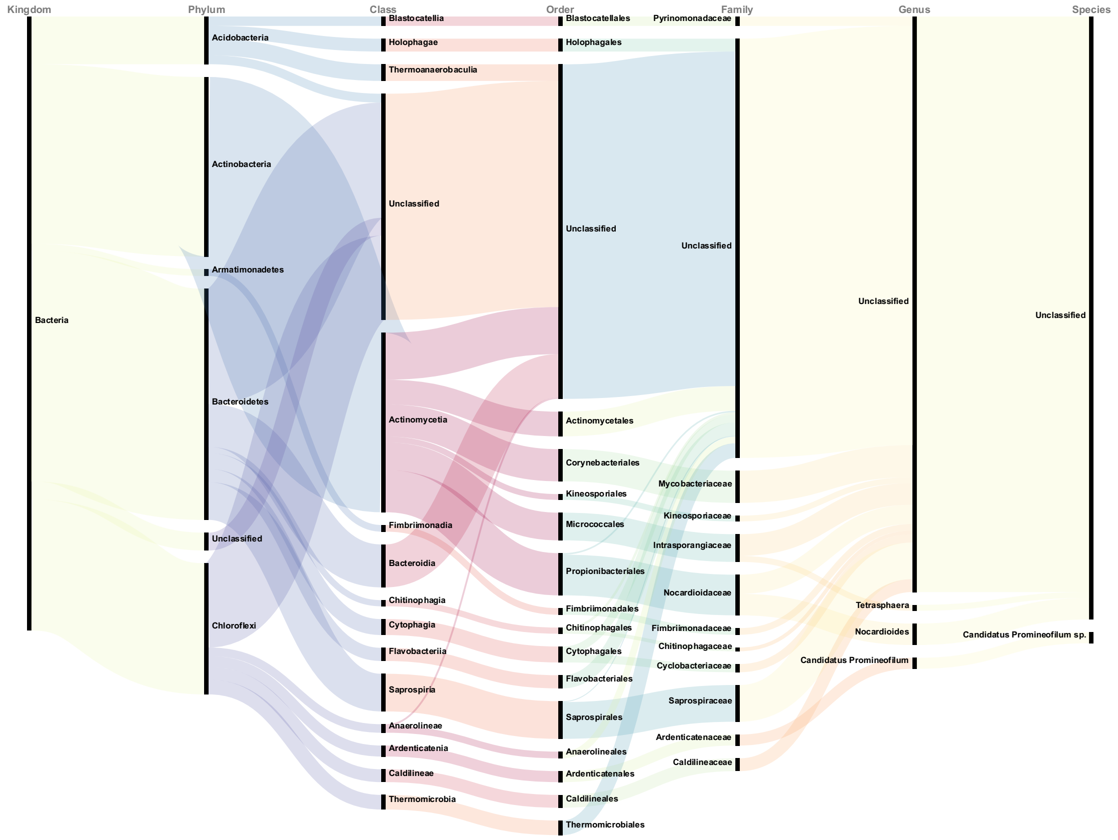

**Figure S23.** Sankey visualization of microbial taxa contributing to K00318 (proline dehydrogenase) under CA conditions.

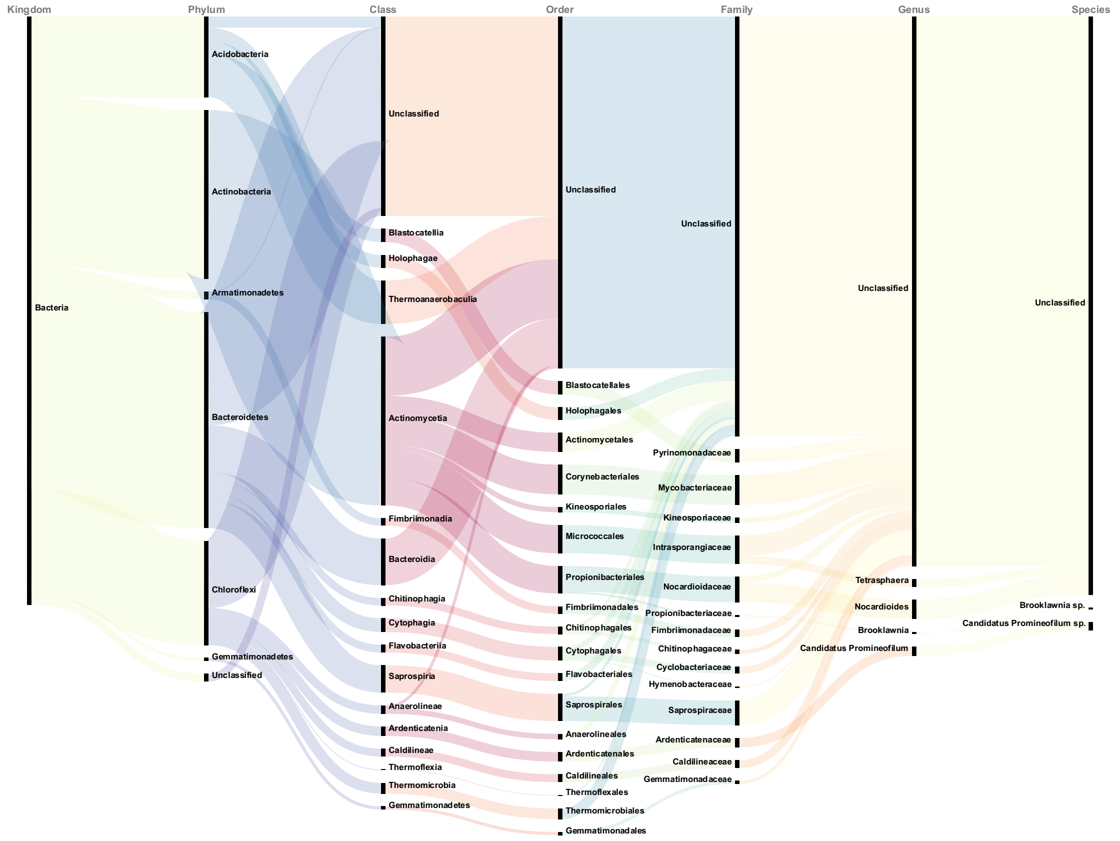

**Figure S24.** Sankey visualization of microbial taxa contributing to K00318 (proline dehydrogenase) under IP conditions.

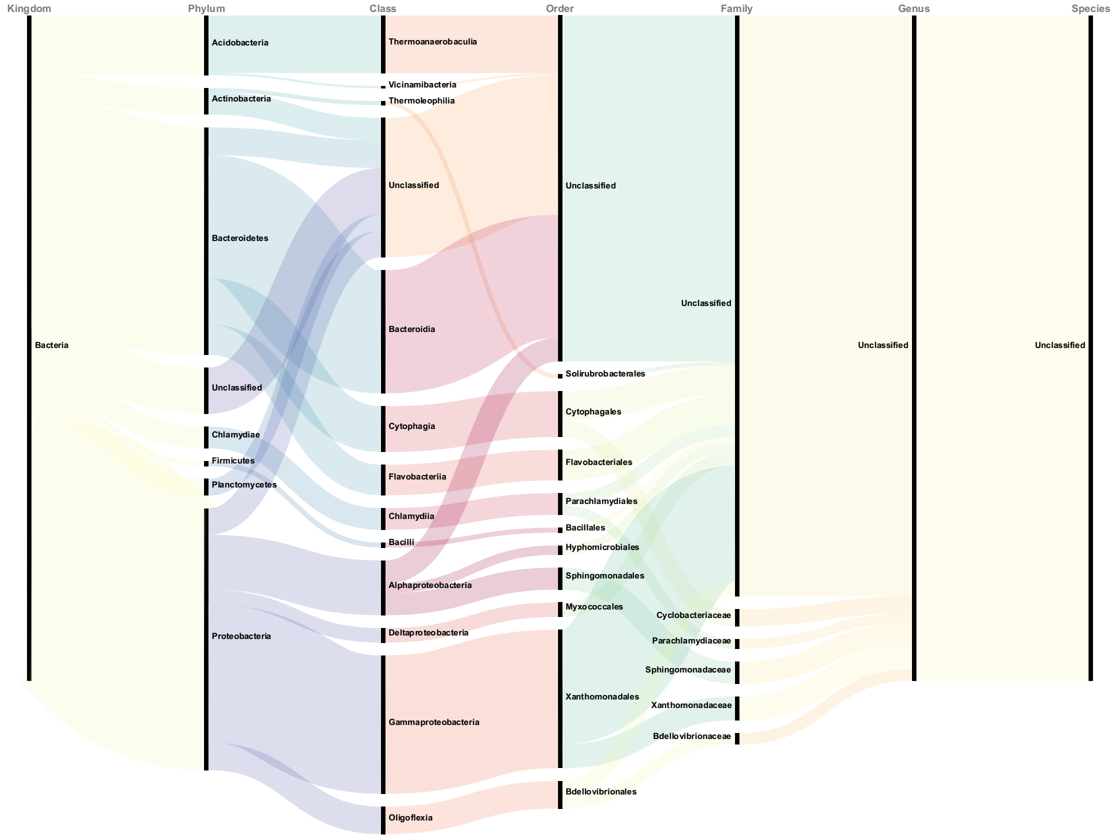

**Figure S25.** Sankey visualization of microbial taxa contributing to K00263 (leucine dehydrogenase) under CP conditions.

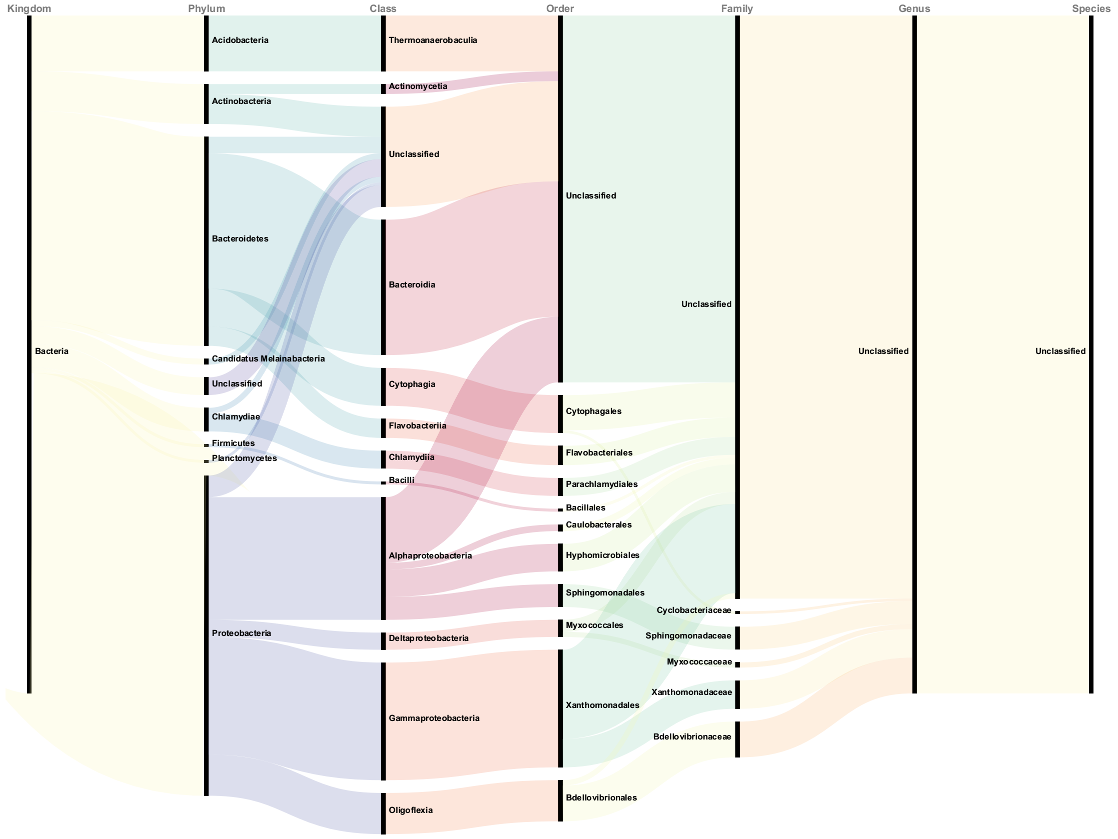

**Figure S26.** Sankey visualization of microbial taxa contributing to K00263 (leucine dehydrogenase) under CA conditions.

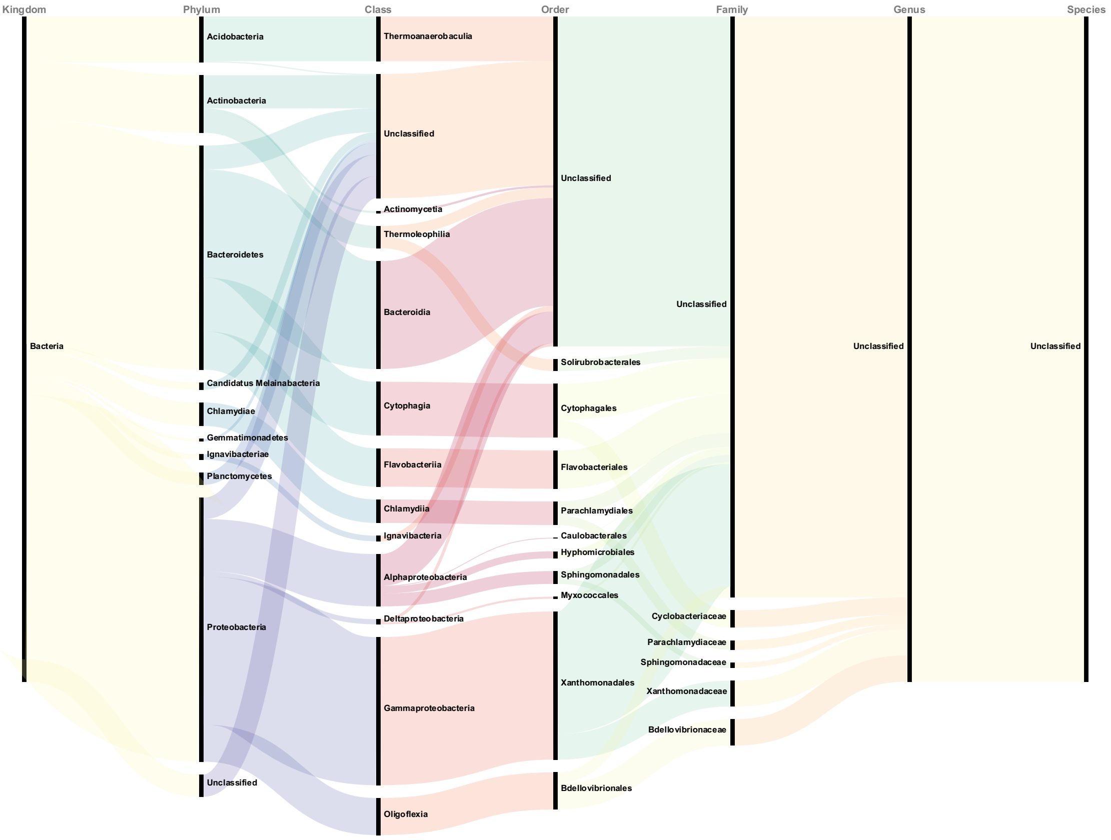

**Figure S27.** Sankey visualization of microbial taxa contributing to K00263 (leucine dehydrogenase) under IP conditions.

**Table S4.** Taxonomic origins of six ROS-generating enzymes identified in the metaproteomic dataset.

| ROS producers | Microbial origins |
|---|---|
| **glutamate synthase (NADPH) large chain [EC:1.4.1.13]** | k_Bacteria;p_Proteobacteria<br>k_Bacteria;p_Proteobacteria;c_Betaproteobacteria;o_Burkholderiales<br>k_Bacteria;p_Proteobacteria;c_Betaproteobacteria;o_Nitrosomonadales;f_Methylophilaceae;g_Methylobacillus<br>k_Bacteria;p_Proteobacteria;c_Alphaproteobacteria;o_Hyphomicrobiales;f_Hyphomicrobiaceae;g_Hyphomicrobium |
| **glutathione reductase (NADPH) [EC:1.8.1.7]** | k_Bacteria;p_Proteobacteria;c_Alphaproteobacteria;o_Hyphomicrobiales;f_Hyphomicrobiaceae;g_Hyphomicrobium |
| **dihydrolipoamide dehydrogenase [EC:1.8.1.4]** | k_Bacteria;p_Proteobacteria;c_Betaproteobacteria<br>k_Bacteria;p_Proteobacteria;c_Betaproteobacteria;o_Burkholderiales |

| | |
|---|---|
| | k_Bacteria;p_Proteobacteria;c_Betaproteobacteria;o_Burkholderiales;f_Burkholderiaceae;g_Burkholderia<br>k_Bacteria;p_Proteobacteria;c_Betaproteobacteria;o_Nitrosomonadales;f_Methylophilaceae<br>k_Bacteria;p_Proteobacteria;c_Betaproteobacteria;o_Nitrosomonadales;f_Sterolibacteriaceae;g_Methyloversatilis<br>k_Bacteria;p_Proteobacteria;c_Alphaproteobacteria;o_Hyphomicrobiales<br>k_Bacteria;p_Proteobacteria;c_Alphaproteobacteria;o_Hyphomicrobiales;f_Hyphomicrobiaceae;g_Hyphomicrobium |
| **glutamate synthase (ferredoxin) [EC:1.4.7.1]** | k_Bacteria;p_Bacteroidetes;c_Saprospiria;o_Saprospirales;f_Saprospiraceae |
| **proline dehydrogenase [EC:1.5.-.-]** | k_Bacteria;p_Actinobacteria;c_Actinomycetia;o_Corynebacteriales;f_Mycobacteriaceae |
| **leucine dehydrogenase [EC:1.4.1.9]** | k_Bacteria;p_Bacteroidetes;c_Bacteroidia |

#### S2.3. Potential yield and economic value of EPS enhancement under full-scale conditions

To evaluate the full-scale applicability of the continuous oxygen perturbation (CP) strategy, we estimated the potential EPS recovery in a municipal wastewater treatment plant (WWTP) assuming an aerobic reactor volume of 3,000 $m^3$, a typical scale for a plant treating 12,000 $m^3$/day of wastewater with a hydraulic retention time of approximately 6 hours.
Based on laboratory measurements, CP resulted in a net EPS increase of 74.3 mg/L/day, while conventional aeration (CA) produced only 39.0 mg/L/day. When scaled to the full aerobic volume:

- CP condition:
  $74.3\ \text{mg/L/day} \times 3{,}000\ \text{m}^3 \times 1{,}000\ \text{L/m}^3 \times 10^{-6}\ \text{kg/mg} = 222.9\ \text{kg/day}$
  Annual EPS yield = 222.9 kg/day × 365 days = 81.4 tons/year
- CA condition:
  $39.0\ \text{mg/L/day} \times 3{,}000\ \text{m}^3 = 117.0\ \text{kg/day}$
  Annual EPS yield = 117.0 kg/day × 365 = 42.7 tons/year

Therefore, the CP strategy could generate an additional 38.7 tons of EPS per year compared to conventional aeration.
Assuming a conservative market value of $5 per kilogram—reflecting the potential application of sludge-derived EPS as low-cost bioflocculants or adsorbents—this additional EPS could yield approximately $193,500 USD in annual economic value. This estimation

does not include potential savings from improved sludge settling or downstream resource recovery, indicating further value may be realized through integrated process optimization.

### References

Bolger, A.M., Lohse, M., Usadel, B., 2014. Trimmomatic: A flexible trimmer for Illumina sequence data. Bioinformatics 30, 2114–2120. https://doi.org/10.1093/bioinformatics/btu170

Buchfink, B., Xie, C., Huson, D.H., 2015. Fast and sensitive protein alignment using DIAMOND. Nat Methods 12, 59–60. https://doi.org/10.1038/nmeth.3176

Guo, G., McKenzie, E.J., Jones, M.B., Zarate, E., Seymour, J. de, Baker, P.N., Villas-Bôas, S.G., Han, T.-L., 2021. MassOmics: An R package of a cross-platform data processing pipeline for large-scale GC-MS untargeted metabolomics datasets. https://doi.org/10.5281/zenodo.4961895

Haug, K., Cochrane, K., Nainala, V.C., Williams, M., Chang, J., Jayaseelan, K.V., O'Donovan, C., 2020. MetaboLights: A resource evolving in response to the needs of its scientific community. Nucleic Acids Research 48, D440–D444. https://doi.org/10.1093/nar/gkz1019

Hyatt, D., Chen, G.-L., LoCascio, P.F., Land, M.L., Larimer, F.W., Hauser, L.J., 2010. Prodigal: Prokaryotic gene recognition and translation initiation site identification. BMC Bioinformatics 11, 119. https://doi.org/10.1186/1471-2105-11-119

Kanehisa, M., Furumichi, M., Sato, Y., Matsuura, Y., Ishiguro-Watanabe, M., 2025. KEGG: biological systems database as a model of the real world. Nucleic Acids Research 53, D672–D677. https://doi.org/10.1093/nar/gkae909

Langmead, B., Salzberg, S.L., 2012. Fast gapped-read alignment with Bowtie 2. Nat Methods 9, 357–359. https://doi.org/10.1038/nmeth.1923

Li, D., Liu, C.-M., Luo, R., Sadakane, K., Lam, T.-W., 2015. MEGAHIT: An ultra-fast single-node solution for large and complex metagenomics assembly via succinct de Bruijn graph. Bioinformatics 31, 1674–1676. https://doi.org/10.1093/bioinformatics/btv033

Muth, T., Behne, A., Heyer, R., Kohrs, F., Benndorf, D., Hoffmann, M., Lehtevä, M., Reichl, U., Martens, L., Rapp, E., 2015. The MetaProteomeAnalyzer: A powerful open-source software suite for metaproteomics data analysis and interpretation. J. Proteome Res. 14, 1557–1565. https://doi.org/10.1021/pr501246w

Perez-Riverol, Y., Bai, J., Bandla, C., García-Seisdedos, D., Hewapathirana, S., Kamatchinathan, S., Kundu, D.J., Prakash, A., Frericks-Zipper, A., Eisenacher, M., Walzer, M., Wang, S., Brazma, A., Vizcaíno, J.A., 2022. The PRIDE database resources in 2022: A hub for mass spectrometry-based proteomics evidences. Nucleic Acids Research 50, D543–D552. https://doi.org/10.1093/nar/gkab1038

Puente-Sánchez, F., García-García, N., Tamames, J., 2020. SQMtools: Automated processing and visual analysis of 'omics data with R and anvi'o. BMC Bioinformatics 21, 358. https://doi.org/10.1186/s12859-020-03703-2

Schmieder, R., Edwards, R., 2011. Quality control and preprocessing of metagenomic datasets. Bioinformatics 27, 863–864. https://doi.org/10.1093/bioinformatics/btr026

Seemann, T., 2014. Prokka: rapid prokaryotic genome annotation. Bioinformatics 30, 2068–2069. https://doi.org/10.1093/bioinformatics/btu153

Smart, K.F., Aggio, R.B.M., Van Houtte, J.R., Villas-Bôas, S.G., 2010. Analytical platform for metabolome analysis of microbial cells using methyl chloroformate derivatization followed by gas chromatography-mass spectrometry. Nat Protoc 5, 1709–1729. https://doi.org/10.1038/nprot.2010.108

Tamames, J., Puente-Sánchez, F., 2019. SqueezeMeta, a highly portable, fully automatic metagenomic analysis pipeline. Frontiers in Microbiology 9.

The UniProt Consortium, 2025. UniProt: The universal protein knowledgebase in 2025. Nucleic Acids Research 53, D609–D617. https://doi.org/10.1093/nar/gkae1010

Xu, M., Li, Z., Li, L., 2013. Combining percolator with X!Tandem for accurate and sensitive peptide identification. J Proteome Res 12, 3026–3033. https://doi.org/10.1021/pr4001256
